# High-throughput formulation of reproducible 3D cancer microenvironments for drug testing in myelogenous leukemia

**DOI:** 10.1101/2024.01.10.575000

**Authors:** M. Rudzinska-Radecka, L. Turos-Korgul, D. Mukherjee, P. Podszywalow-Bartnicka, K. Piwocka, J. Guzowski

## Abstract

Targeting cancer microenvironment is currently one of the major directions in drug development and preclinical studies in leukemia. Despite the variety of available chronic myelogenous leukemia 3D culture models, the reproducible generation of miniaturized leukemia microenvironments, suitable for high-throughput drug testing, has remained a challenge. Here, we use microfluidics to generate over ten thousand highly monodisperse leukemic-bone marrow hydrogel microbeads per minute. We employ gelatin methacrylate (GelMA) as a model extracellular matrix (ECM) and tune the concentration of the biopolymer, as well as other possible components of the ECM (fibrin, hyaluronic acid), cell concentration and the ratio of leukemic cells to bone marrow cells within the microbeads. This allows to achieve optimal cell viability and the propensity of the encapsulated cells to microtissue formation, while also warranting long-term stability of the microbeads in culture. We administer model kinase inhibitor, imatinib, at various concentrations to the microbeads and, via comparing mono-and co-culture conditions (cancer alone vs cancer-stroma), we find that the stroma-leukemia crosstalk systematically protects the encapsulated cells against the drug-induced cytotoxicity, confirming therefore that our system mimics the physiological stroma-dependent protection. We finally discuss applicability of our model to (i) studying the role of direct-or close-contact interactions between leukemia and bone marrow cells embedded in 3D ECM on the stroma-mediated protection, and (ii) high-throughput screening of anti-cancer therapeutics in personalized therapies.

## Introduction

The development of cancer microenvironment (CME) models has become one of the critical objectives in preclinical studies. CMEs have been previously addressed mostly in the context of solid tumors, however, more recently, it has become evident that the proper modeling of the microenvironment in cancers not forming solid tumors, such as myeloid leukemias, including chronic myeloid leukemia (CML), remains equally important (Federica Barbaglio et al., 2020; Hermansen et al., 2023; Scielzo and Ghia, 2020).

Unfortunately, the physiologically relevant and reproducible leukemia CME models are lacking. While the standard suspension leukemic cell cultures are not fully suitable to address the interactions of the leukemic cells with the bone marrow microenvironment (BMM), the available *in vivo* or *ex vivo* models – even though physiologically relevant – have significant limitations including, among others, high costs, low throughputs (as animal experiments are time-consuming), and low reproducibility. At the same time, the worldwide trends in biomedical research as well as the new FDA and EU regulations are inevitably directed toward the reduction of animal testing and the development of more reliable preclinical research methods (Moutinho, 2023). In particular, the new techniques of 3D cell culture capable of recapitulating the biological complexity of the leukemic CME via combining multiple CME-associated cell types together with the cancer-specific extracellular matrix (ECM) in a predefined (biomimetic) spatial arrangement are urgently needed.

In general, the development of leukemic 3D microenvironment models remains challenging due to the intrinsic complexity of the leukemic CME which includes a variety of bone marrow cells and signaling pathways (Simioni et al., 2021). For example, it has been previously shown that the myeloid malignant cells can induce extensive niche remodeling when growing in the bone marrow and disrupt the normal perivascular niche by reducing CXCL12 production (Colmone et al., 2008). Furthermore, cancer cells produce large amounts of stem-cell factor (SCF), attract normal hematopoietic stem cells and modify their proper functioning (Duan et al., 2014). Additionally, as we have shown in our previous work, leukemic cells with an active-adaptative stress response, secrete enzymes that modulate the extracellular matrix and remodel invasiveness potential of the cells (Podszywalow-Bartnicka et al., 2016). On the other hand, stromal cells, by either direct-or indirect crosstalk with leukemic cells, significantly change their biology by activating a protective response. As a result — via enhancing the metabolic plasticity, pro-survival signaling and proliferation of leukemic cells — the bone marrow cells may favor the persistence and progression of leukemia, including the self-renewal capacities of the malignant progenitor cells (Vianello et al., 2010). Overall, the bone marrow microenvironment is known to contribute to myeloid malignancies and protect CML cells from drug-induced cytotoxicity through various routes of intercellular communication (Chen et al., 2021; Dolinska et al., 2023; Joshi et al., 2021; Kolba et al., 2019; Korn and Méndez-Ferrer, 2017; Le et al., 2020; Pal et al., 2022; Patterson and Copland, 2023). For example, immunosuppression and T-cell exhaustion (Swatler et al., 2022b, 2022a) are promoted in the presence of leukemic extracellular vesicles (EVs).

Importantly, stromal cells serve as the driver of cytoprotective signaling in leukemia, participating in developing resistance to anti-cancer treatments. This includes resistance to tyrosine kinase inhibitors (TKIs) (Deininger et al., 1997; Druker et al., 2001, 1996; Melo and Chuah, 2007; Kolba et al., 2019; Kumar et al., 2017) as well as against PARP1 inhibitors (Le et al., 2020; Podszywalow-Bartnicka et al., 2019; Tobin et al., 2013) considered the novel personalized therapy in leukemias. Various types of interactions between the bone marrow stromal and leukemic cells that influence therapeutic effectivity in myeloid leukemias have been reported (Park et al., 2022; Saito et al., 2021; Kolba et al., 2019; Le et al., 2020; Wolczyk et al., 2023), therefore they should be addressed when designing the effective therapeutic strategy. Furthermore, it has been shown by others (Schepers et al., 2013) that leukemic cells can actively remodel their environment, and such bone marrow niche dysregulation negatively affects treatment outcomes (Agarwal et al., 2017; Zhang et al., 2016, 2013). Overall, targeting the elements of the bone marrow environment is currently considered one of the promising directions toward successful therapies for leukemia that alleviate the acquisition of a chemoresistant phenotype. Therefore, the drug development process should also address the microenvironment effects.

Most of the previous approaches toward modeling drug resistance in the leukemic CME have been limited to 2D cultures or the suspension co-cultures of leukemic and bone marrow cells. Despite the superior physiological relevance of the 3D culture, reliable 3D leukemia CME models that could mimic the close interactions of cancer-stromal cells, the adhesion of cells to the extracellular matrix, and the signaling of soluble factors that modulate their response to therapeutics (Al-Kaabneh et al., 2022; Araujo-Ayala et al., 2023; Fontana et al., 2021; Habanjar et al., 2021; Scielzo and Ghia, 2020)) remain scarce. Here, we use droplet microfluidics to co-encapsulate human CML cancer cells (K562) and stromal bone marrow cells (HS-5) inside gelatin methacrylate (GelMA) hydrogel microbeads and we demonstrate their use as miniature, highly reproducible models of the leukemic bone marrow microenvironment and leukemia-stroma crosstalk.

In particular, we use our leukemic microenvironment model to study the stroma-mediated protection of the leukemic cells against a model drug (tyrosine kinase inhibitor - imatinib), which remains a major challenge in the effective eradication of leukemic cells. In fact, despite excellent efficacy and improved clinical response levels acquired by tyrosine kinase inhibitors in leukemic patients in the chronic phase, the effects of drug resistance or relapse are observed in a significant number of patients, even after the initial success (Bhamidipati et al., 2013; Jabbour et al., 2013). This resistance is an emerging problem in clinical practice, and a challenge in CML research and drug discovery (Jabbour et al., 2011)). The precise and reproducible leukemic microenvironment models that we demonstrate in this work may be utilized directly in leukemia research or developed further towards an integrated platform incorporating automated methods of microtissue manipulation, e.g., via their bioprinting (Germain et al., 2022; Wei et al., 2022). Such platforms could be used for high-throughput studies on the mechanisms of drug resistance and other pre-clinical tests on leukemic cells as well as for the development of personalized anti-cancer therapies.

## Results

In our leukemic bone marrow microenvironment model, we use the BCR-ABL1-positive chronic myeloid leukemia (CML) K562 cells as the leukemic cell line and HS-5 as the stromal cell line. The HS-5 cell line can reproduce the effect of primary bone marrow MSCs in tumor biology and immunomodulation and therefore it has been widely utilized as a model bone marrow stromal cell type in leukemia studies (Adamo et al., 2020). Both cell lines are commonly used as model cells to study leukemia bone marrow microenvironment as well as for drug screening. As a model drug we use Imatinib (STI571), a first-generation tyrosine kinase inhibitor (TKI) approved for frontline therapy in CML patients by the US Food and Drug Administration (FDA) in 2002 (Bhamidipati et al., 2013). By attaching closely to the ATP binding site of BCR-ABL, imatinib stabilizes the inactive conformation of BCR-ABL1, thus inhibiting the BCR-ABL-specific tyrosine kinase activity (Hantschel et al., 2012). Resistance to imatinib appears at later stages of the disease due to internal or external (microenvironment-dependent) pro-survival signaling. In our experiments, we co-encapsulate the mixture of K562 and HS-5 cells inside GelMA-based hydrogel microbeads, culture the microbeads for up to 72 h, and screen the viability of the encapsulated microtissues against different concentrations of Imatinib in the bone marrow-mimicking conditions.

## 1. Optimization of hydrogel composition in terms of finest cell growth

GelMA is a methacrylated derivative of gelatin and, similar to collagen, contains the tripeptide Arg-Gly-Asp sequence for cell adhesion and the matrix metalloprotease (MMP) degradation sequence, the latter allowing for matrix remodeling (Wang et al., 2016). In addition, GelMA undergoes fast and efficient photo-crosslinking upon exposure to UV light in the presence of a photoinitiator which makes it suitable for a variety of tissue-engineering applications (Nichol et al., 2010).

Mechanical properties of GelMA hydrogels can be easily tuned by varying GelMA concentration, degree of methacrylation, the concentration of photoinitiator, and UV irradiance (Im and Lin, 2022; Liu et al., 2012; Velasco-Rodriguez et al., 2021; Wang et al., 2014). It is well known that, in general, the type of biopolymer forming the hydrogel matrix, as well as the stiffness of the crosslinked hydrogel matrix can influence the viability, proliferation, and differentiation of the embedded cells (Klotz et al., 2016; Zhao et al., 2016a). In particular, it is also known that hybrid hydrogels composed of GelMA and methacrylated derivative of hyaluronic acid (HAMA) tend to be more stable in culture, i.e., have slower degradation rates. However, the addition of HAMA also leads to matices of higher stiffness which typically result in slower cell proliferation and delayed microtissue formation (Camci-Unal et al., 2013). On the other hand, the use of much softer hydrogels such as fibrin has been shown to promote cell migration and differentiation, however, the efficiency of GelMA photo-crosslinking in the presence of fibrin and its impact on CML cell viability remain unclear. Therefore, we tested the impact of HAMA and fibrin addition to the GelMA matrices in terms of leukemic and bone-marrow cell proliferation. To this end, we performed CCK-8 assay and confocal 3D live/dead imaging of K562 and HS-5 cells embedded inside GelMA, GelMA/HAMA, or GelMA/fibrin hydrogel slabs of thickness around 1 mm (Figure 1). We tested the following hydrogel samples: 3.2% GelMA, 4.8% GelMA, 4.8% GelMA with 0.4% HAMA, 4.8% GelMA with 0.08% Fibrin and 4.8% GelMA with 0.2% Fibrin. Considering cell viability, we observed the most active cell proliferation in 4.8% GelMA, and 4.8% GelMA + 0.4% HAMA, with no significant difference between these two variants (Figure 1A). The addition of fibrin caused excessive cell sedimentation, visible as cells growing mostly at the bottom of the dish, rather than within the hydrogel. The same effect, yet less pronounced, was observed in the case of 3.2% GelMA (Figure 1B). Moreover, a large fraction of dead cells was observed in the samples with added fibrin. This impaired viability might be related to the non-full crosslinking of the hydrogel and the presence of free radicals formed upon 405 nm irradiance. Such behaviour excludes the possibility of long-term culture inside microgels, therefore in the further microfluidic encapsulation experiments we only used the variants without fibrin, that is 4.8% GelMA and 4.8% GelMA + 0.4% HAMA.

**Figure 1.**
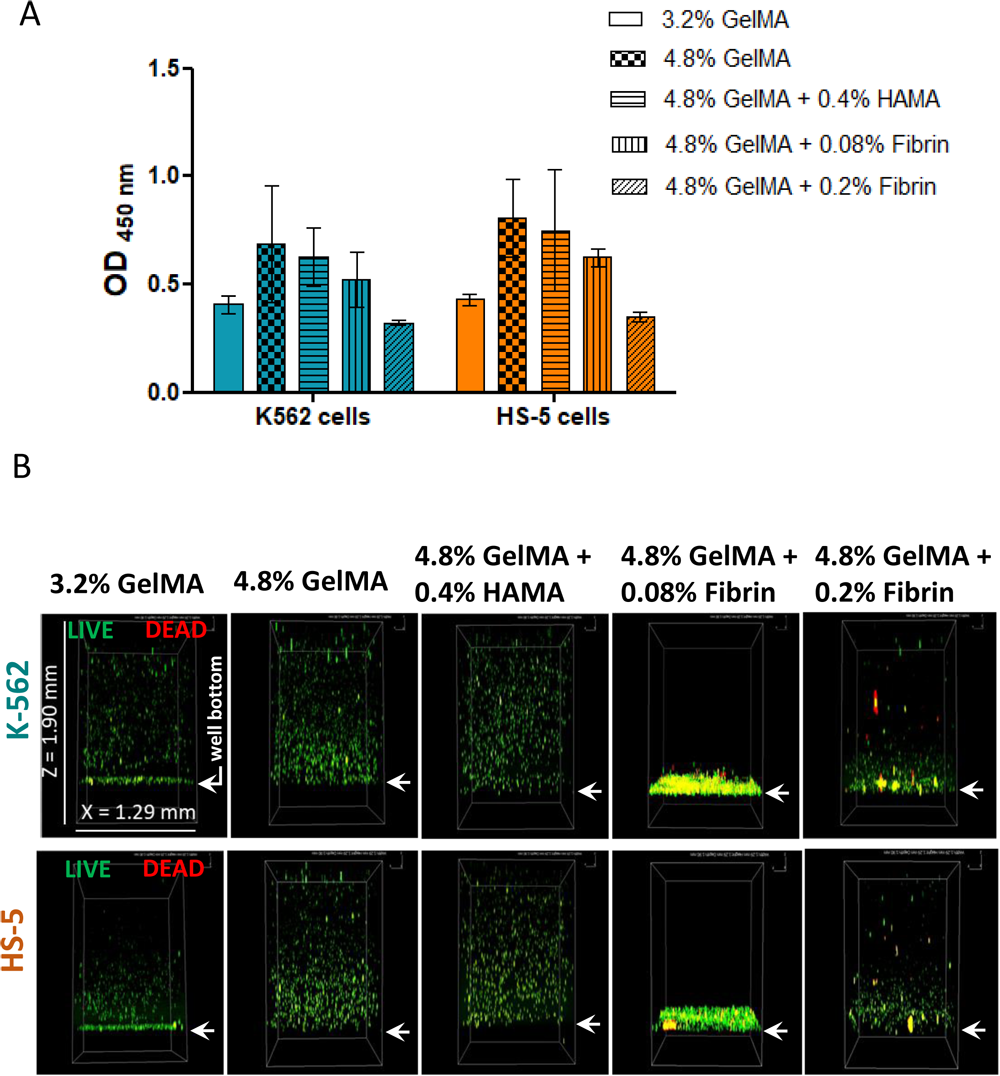
Cell proliferation evaluated using **(A)** CCK-8 assay (absorbance at 450 nm) after 48 h of culture for cells evenly mixed within the hydrogel; **(B)** 3D confocal microscopy profiles of cells distributed in the hydrogel layer (∼1.6 mm thickness) in a 96-well plate after 48h of culture showing viable (green) and non-viable (red) cells. Arrows indicate well bottom.

## 2. Encapsulation of chronic myeloid leukemia and human bone marrow stromal cells using microfluidic devices

It is known that the small dimensions of the microbeads generated with microfluidics allow for efficient diffusion of oxygen, exchange of metabolites, and delivery of nutrients to the encapsulated cells (Sart et al., 2022), thus supporting proliferation and ultimately formation of dense tumor-stroma microtissues (Rojek et al., 2022). The use of microgels as microtissue carriers additionally facilitates manipulation and imaging of the individual microtissues while also providing a physical barrier that protects the cells from external mechanical stresses.

In our experiments, we encapsulated K562 and HS-5 cells within hydrogel microdroplets using a microfluidic device which generated droplets at a frequency of around 12 000/min. The device consisted of an inlet for the hydrogel phase with suspended cells, another inlet for the oil phase, and a single outlet for the generated emulsion. Both hydrogel and the oil phases were supplied at constant rates of flow. The hydrogel phase was distributed into multiple independent droplet generators which created droplets via the so-called step-emulsification (Sugiura et al., 2001), see Figure 2A. The droplets were then carried away by the oil phase and collected outside the device. The advantage of the step-emulsification device over other methods of droplet generation (flow-focusing, T-junction, etc.) is that the size of the droplets is predominantly set by the channel geometry and only weakly depends on the applied rates of flow. Step-emulsification is particularly advantageous in the case of high-throughput generation of droplets in multiple parallel junctions supplied from a single source (De Rutte et al., 2019). Due to the hydrodynamic crosstalk between the different junctions, the actual rates of flow in the individual junctions naturally fluctuate over time. However, in the case of step-emulsifiers such fluctuations have little effect on the size of the generated droplets. Our device generated droplets of mean diameter *D* = 202 (±8) µm with the coefficient of variation CV = 3.8 % (Figure 2B) under the applied rates of flow equal Q_hyd_ = 50 µl/min for the cell-hydrogel mixture and Q_oil_ = 150 µl/min for the oil phase (Novec 7500-surfactant mixture). We therefore expect an average of 33.5 cells/droplet in the case of cell concentration 8 x 10^6^/ml. From direct counting, we obtained typically between 9 and 30 cells per droplet with an average of 15.6 cells/droplet and CV = 37 %. This suggests some degree of cell clustering such that pairs or groups of cells appear as a single object. In fact, in the case of the absence of any direct cell-cell interactions (no clustering), the distribution of cells in droplets would follow the Poisson distribution (dotted line in Figure 2C). However, we find a better fit with a slightly wider Gaussian distribution (solid line in Figure 2C) which remains in line with the picture of the moderate pairing/clustering.

**Figure 2.**
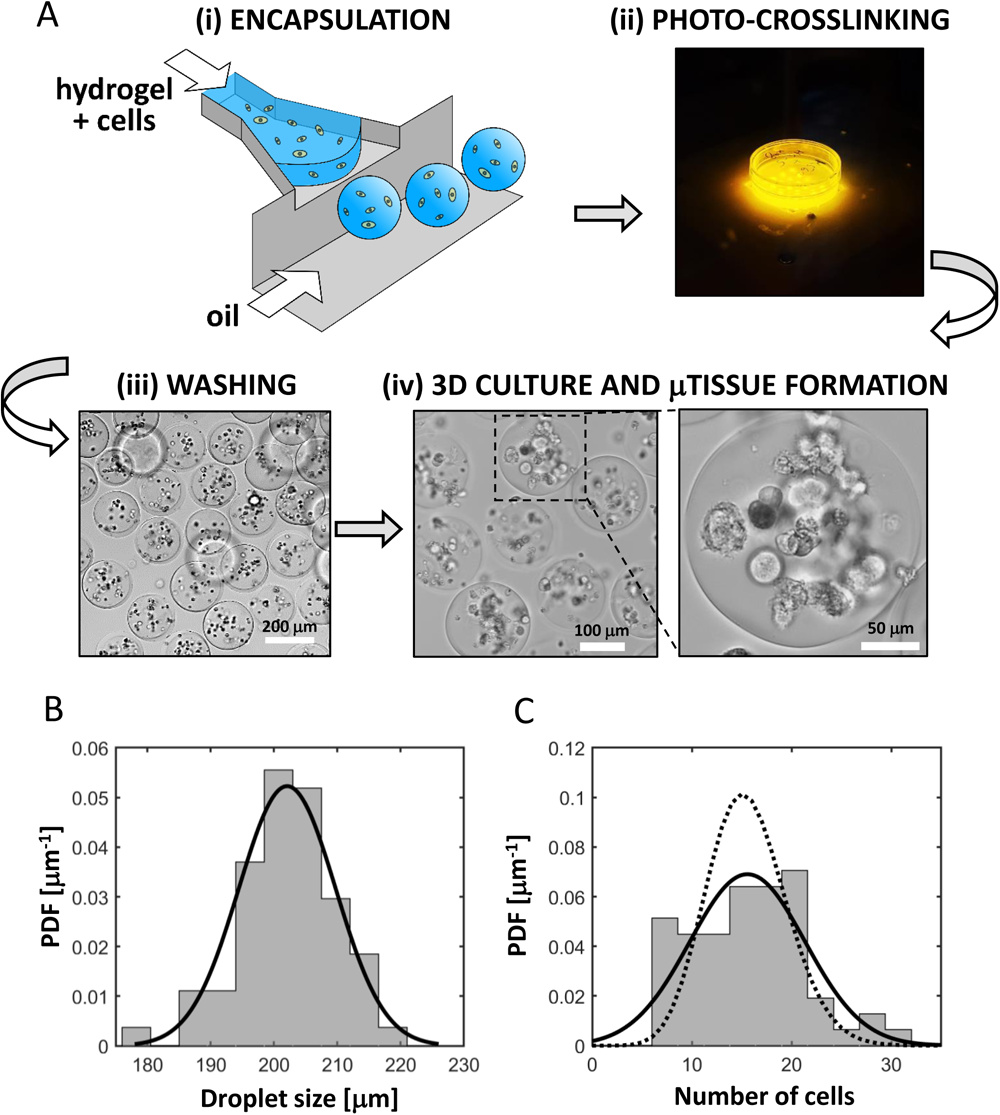
Cell encapsulation in droplet microfluidic platform **(A)**: (i) generation of microdroplets encapsulating multiple cells using a step-emulsification device, (ii) photocrosslinking with a custom 405 nm LED array, (iii) washing out the oil, (iv) long-term culture of cell-laden microbeads; **(B)** Droplet size distribution fitted with a Gaussian curve (solid line); **(C)** Distribution of the number of cells encapsulated per droplet fitted with a Gaussian curve (solid line) and a Poisson curve (dotted line).

The microdroplets with encapsulated cells were collected and crosslinked using the custom-fabricated 405 nm LED array for 10 sec under the irradiance of 280 mW/cm^2^. The cell-encapsulated microbeads were washed and maintained in culture for 72 h and observed under the microscope.

We tested two different biomaterials, 4.8% GelMA and 4.8% GelMA + 0.4% HAMA using cell/hydrogel mixtures to produce the microbeads. First, we compared the growth of both cell lines K562 or HS-5, after 72 h of culture inside the microbeads at cell concentration 8 x 10^6^/ml (Figure 3A). We observed better cell growth and faster spheroid formation in the case of 4.8% GelMA, especially in the case of K562 cells. Therefore, we thereafter used 4.8% GelMA and optimized cell concentration in the hydrogel. To this end, we additionally tested the cell concentrations 6 x 10^6^/ml and 12 x 10^6^/ml (Figure 3A). We observed that the number of spheroids inside the beads gradually increased with the cell concentration (Figure 3 B), from around the average of 2.5 or 1.5 spheroids per bead at 6 x 10^6^ cells/ml to the maximum of around 9 or 6 spheroids per bead at 12 x 10^6^/ml and, for K562 or HS-5 cells, respectively. However, in the case of 12 x 10^6^ cells/ml, we observed occasional cell outmigration and colonization of the substrate (bottom of the well plate, Figure 3D) which indicated that higher cell concentrations may be not suitable for long-term culture, especially beyond 72 h. Therefore, in most further experiments (i.e., in the co-culture ratio assays, drug testing assays, Figs. 5-7) we used the concentration 8 x 10^6^/ml. For comparison of monoculture vs co-culture growth rate (Fig. 4) we used 12 x 10^6^/ml cells in total.

**Figure 3.**
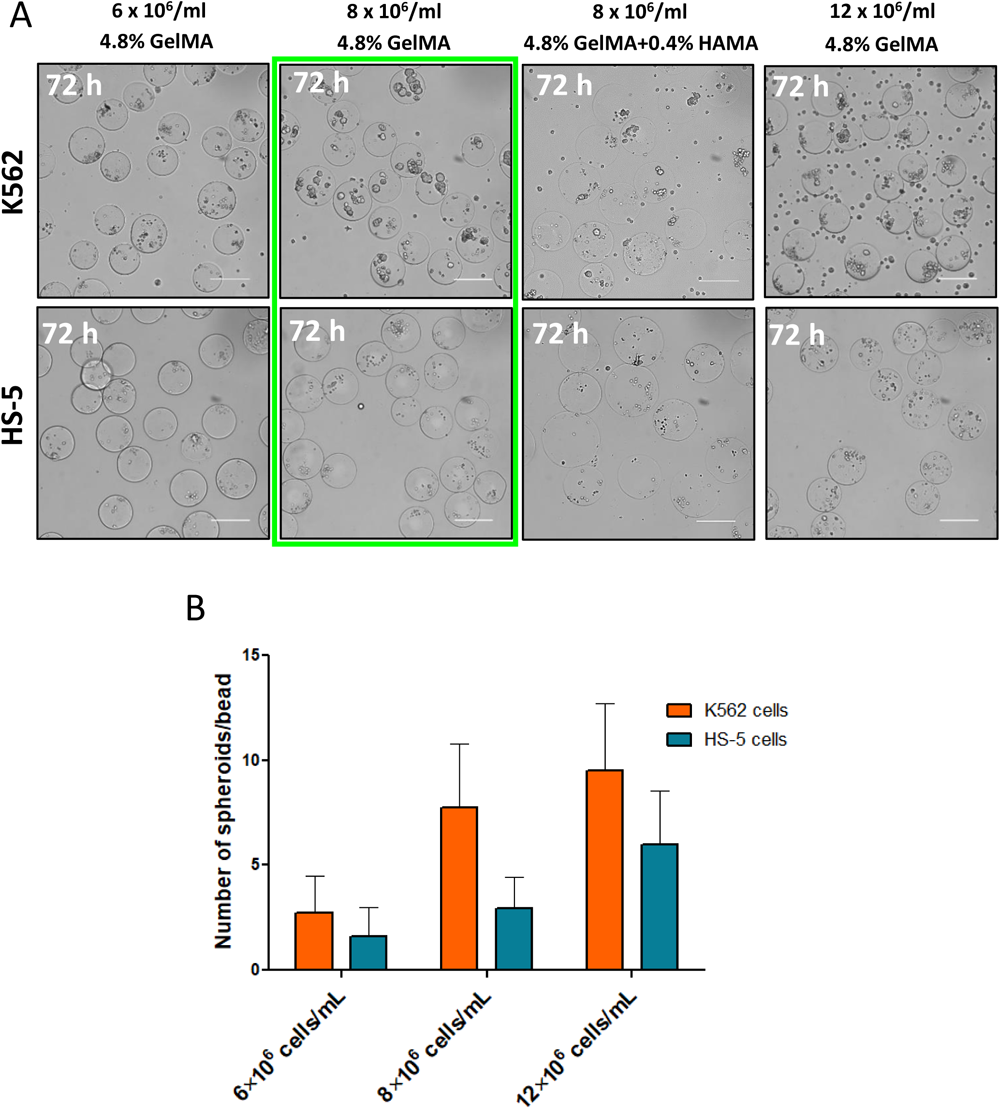
**(A)** Brightfield images of 4.8% GelMA microbeads with encapsulated K562 or HS-5 monocultures at 72 h post encapsulation for various cell concentrations (6 x 10^6^/ml, 8 x 10^6^/ml, 12 x 10^6^/ml). At the cell concentration 8 x 10^6^/ml we additionally show the case with 4.8 % GelMA + 0.4 % HAMA as the hydrogel. Scale bar is 200 µm. The green frame indicates the optimal condition with good cell growth and limited cell outmigration. **(B)** Counted number of spheroids per microbead for 4.8% GelMA microbeads at different cell concentrations for both K562 and HS-5 monocultures.

**Figure 4.**
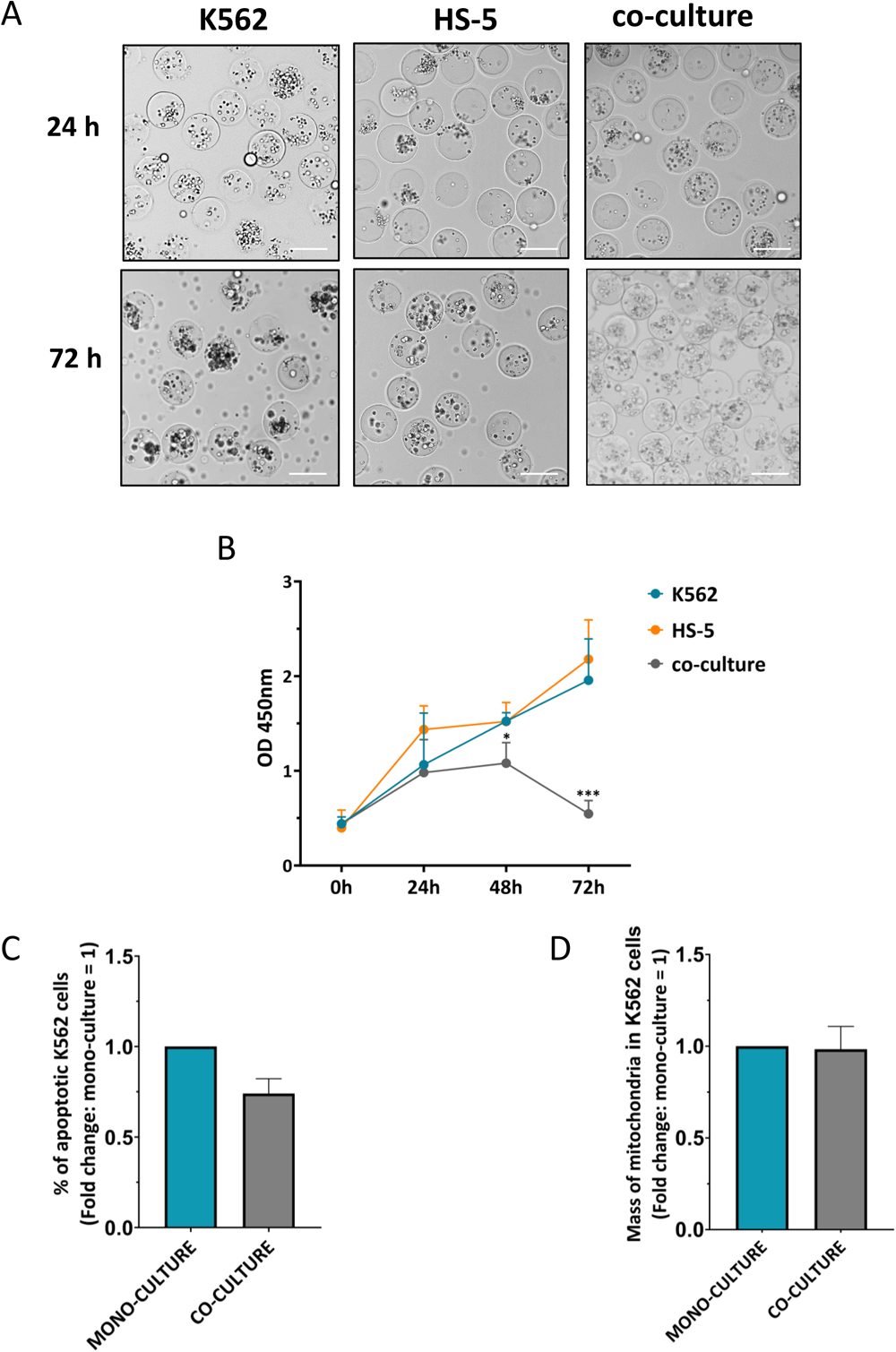
Monoculcure vs co-culture (1:1; K562:HS-5) of leukemia and stroma cells in 4.8% GelMA microbeads. **(A)** K562 and/or HS-5 cells encapsulated and cultured for 72 h. **(B)** Proliferation measured using CCK-8 assay. Cell growth in the co-culture conditions is significantly decreased compared to HS-5 and K562 monocultures; \**p* < 0.05, \*\**p* < 0.01; Mann-Whitney test; results are expressed as mean ± SD. **(C)** Apoptosis and **(D)** mitochondrial activity of K562 cells, measured by flow cytometry in cells cultured in microbeads in co-culture versus monoculture conditions at 72h post encapsulation. In all cases cell concentration in microbeads was 12 x 10^6^ cells/ml.

**Figure 5.**
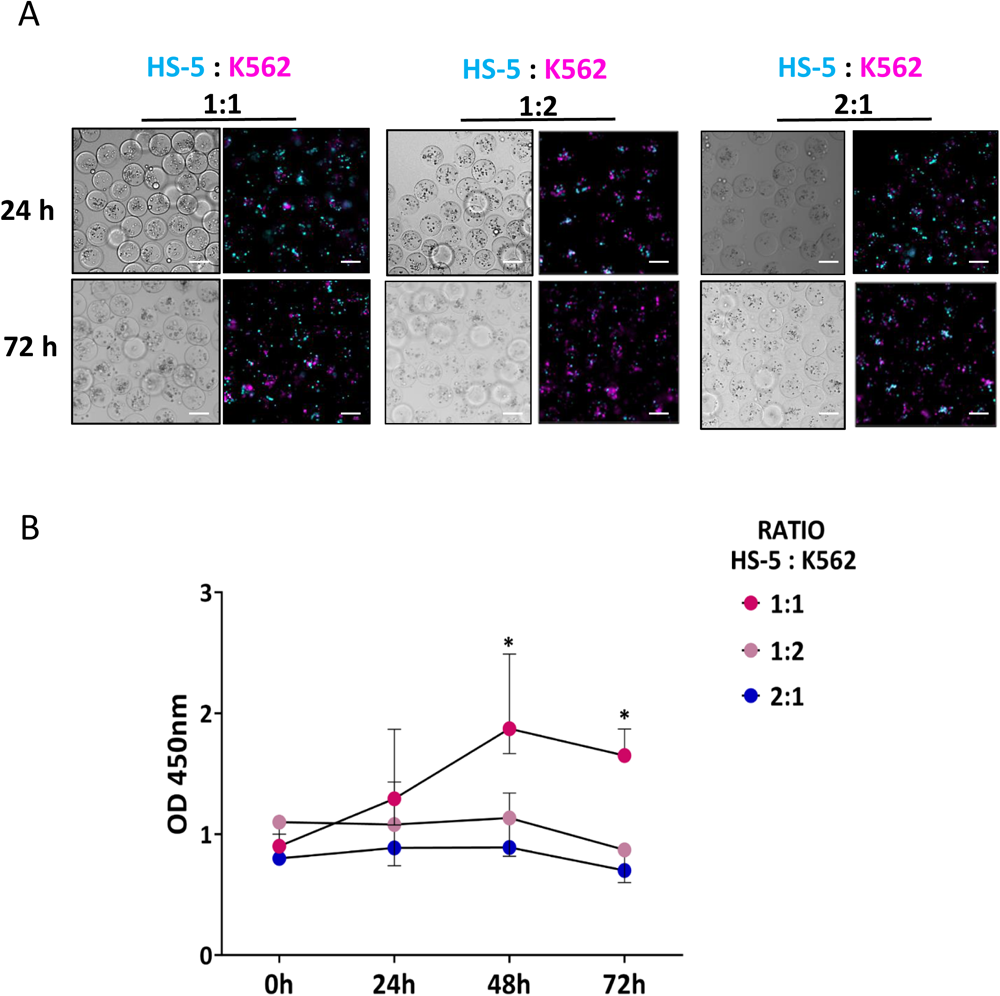
Co-culture of leukemia (magenta) and stroma (cyan) cells (artificially colored, original labels: K562 -red CMTPX and HS-5 – green CMFDA) at various ratios inside microbeads. **(A)** K562 and/or HS-5 (total cell conncentration = 8 x 10^6^/mln) were encapsulated and cultured for 72 h in 1:1, 1:2 or 2:1 ratio of HS-5 : K562. Scale bar = 200 µm. **(B)** Proliferation was measured using CCK-8. Co-culture at 1:1 ratio warrants fastest growth, significantly faster than the 1:2 and 2:1 \**p*> 0.5, Mann-Whitney test; results are expressed as mean ± SD.

## 3. Leukemia-bone marrow interactions within hydrogel microbeads

The inhibition of the bone marrow-derived cell growth by the leukemic accompaniment (leukemia cells and/or conditioned medium) was previously demonstrated by Aoyagi et al., (Aoyagi et al., 1994) using leukemia-derived cell lines with various lineage characteristics and stromal cells (KM-101) or bone marrow culture derived from normal bone marrow. We also investigated the impact of leukemia-stroma interaction in our system.

First, we compared cell growth in the case of the leukemic microenvironment beads, that is with co-cultured (co-encapsulated) K562 and HS-5 cells (ratio 1:1), as well as in the case of the monoculture beads (Figure 4A). The growth of cells in the co-culture measured after 48h was slightly slower than in monocultures. We have previously observed similar effects in 2D co-cultures (Wolczyk et al., 2023). After another day in culture, that is at 72h, we observed a significant drop in the number of proliferating/growing cells (Figure 4B). Therefore, we have selected 48-hour co-culture as an optimal condition for drug testing experiments, not biased by either decreased proliferation or the egression of cells outside the beads.

Flow cytometric analysis of apoptosis and mitochondrial activity, measured selectively on fluorescently tracked and separated cell types, confirmed that leukemic cells encapsulated in co-culture with stromal cells showed good condition and vitality (Figure 4C, D). We even observed lower basal apoptosis in co-cultured leukemic cells, which confirms the pro-survival role of the stromal component.

To further verify our culture conditions, the cells were encapsulated in the beads with different ratios of the stromal cells to the leukemia cells (HS-5:K562), namely 1:1, 1:2, and 2:1, see Figure 5A. To visualize and track HS-5 and K562 cells in co-culture, K562 cells were labeled with CellTracker™ Red CMTPX Dye (re-colored in image post-processing to magenta) and HS-5 with CellTracker™ Green CMFDA Dye (re-colored to cyan), both suitable for long-term cell tracking. After cell labeling and encapsulation, the microbeads were washed, resuspended in a culture medium, and cultured for up to 72 h on 96-well plates (Figure 5A). Based on the fluorescence microscopy images we found that the 1:1 ratio significantly shows the highest viability as quantified using CCK-8 assay (Figure 5B), around 2x higher (*p* < 0.5) as compared to the 1:2 and 2:1 ratio.

## 4. Impact of the microenvironment on imatinib resistance in leukemic microenvironment microbeads

Stromal cells were previously reported to provide chemoprotection to CML cells via secreted factors and direct cell-cell communication (Kolba et al., 2019; Kumar et al., 2017; Podszywalow-Bartnicka et al., 2018; Weisberg et al., 2008; Wolczyk et al., 2023). Various factors secreted by stromal cells, including inflammatory cytokines like IL-6 and IL-8, or TGFβ can protect CML cells from BCR-ABL1 kinase inhibition by kinase inhibitors such as imatinib (Civini et al., 2013; Le et al., 2020).

Previously, we have detected direct intercellular tunneling nanotubes between stromal and cancer cells that mediated the transfer of cellular vesicles and promoted chemoresistance (Kolba et al., 2019). Therefore, here, we also analyzed the possibility of the formation of a direct cellular connection between co-cultured cells (HS-5 cells – cyan and K562 cells – magenta, see Figure 6A). In some cases, we indeed found 3D spatial overlap between fluorescence signals coming from HS-5 and K562 cells. We used Nikon NIS-elements software to measure Pearson’s correlation coefficient *C* for cells within a single bead (Figure 6A) inside several chosen ROIs covering pairs of closely neighboring cells. We found the maximal values in the selected confocal z-slices to be *C* = 0.81, 0.45, 0.43, and 0.36 respectively in ROI 1, ROI 2, ROI 3, and ROI 4 (as indicated in Fig. 6A), and similar *C*-values in the neighboring z-slices. The observed high values of *C*, which is close to or larger than 0.5, confirm significant 3D overlap. These results suggest that the formation of direct interactions between cancer and stromal cells in microbeads is possible. Therefore, in the future, the proposed model could be used also in the analysis of direct cancer-stroma connections.

**Figure 6.**
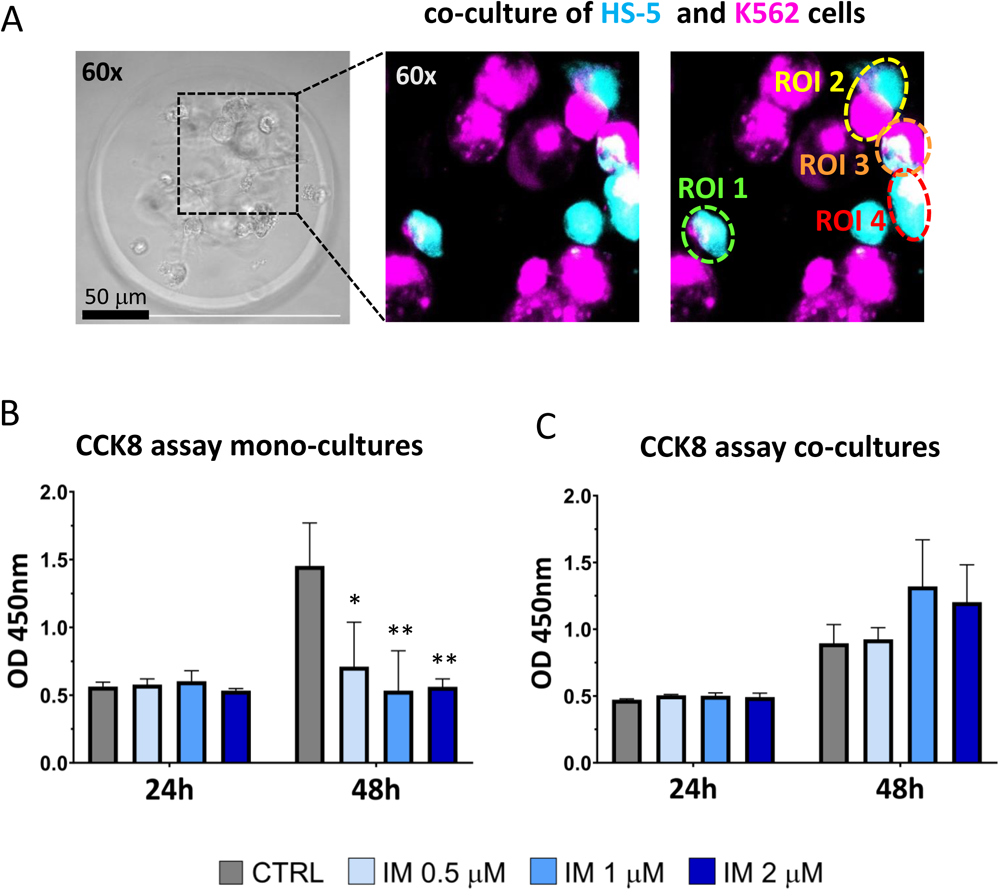
**(A)** The 3D co-culture (K562:HS-5) in 4.8% GelMA microbeads supports formation of direct cellular interactions. Pictures were taken after 48h of culture using confocal microscope. The fluorescent maximum intensity projection reveals close contacts between stroma (cyan) and tumor (magenta) cells with visible overlaps (white). Regions of interest (ROIs 1, 2, 3, 4) identify 4 different pairs of cells with significant 3D overlap measured via Pearson’s coefficient (see main text). Cell proliferation measured using CCK-8 assay for leukemia K562 cells **(B)** in monoculture and **(C)** co-cultured with HS-5 during the continuous treatment with 0.5 µM, 1.0 µM, and 2.0 µM imatinib. In the case of monoculture the increasing drug concentration leads to decreased viability at 48h post encapsulation (i.e., 44h post treatment). On the contrary, in the case of co-culture there is no apparent effect of drug on the growth of cells measured at the same time-point.

Finally, to verify the possibility of the development of drug resistance in our model, the K562 monoculture or the K562: HS-5 (1:1) co-culture (the former as a control) were encapsulated in 4.8% GelMA beads and, after 4h of culture, treated with 0.5-2.0 µM imatinib for the next 72 h (Figure 6B, C). Cytotoxicity assay for drug sensitivity comparison (CCK-8 assay; Figure 6B, C) was assessed at two time points: 24h and 48h. The 72-hour time-point has been excluded from the cytotoxicity drug testing as being highly biased due to the decreased cell growth (Fig. 4B, 5B) as well as possible cell out-migration.

In the leukemic monoculture, imatinib led to a dose-dependent decrease in cell growth compared to the untreated cells, as evidenced by the lower OD 450nm value in the CCK-8 test (Fig. 6B). Additionally, LIVE/DEAD (green/red) fluorescence staining of cancer cells confirmed the imatinib-induced cell death (Supplementary Figure S1). To study the effect of stromal component on sensitivity to imatinib treatment, cells were co-cultured and treated with imatinib at the same doses as in the case of monoculture. In the co-culture conditions, we observed significant protection of the leukemic cells from drug cytotoxicity. In fact, the growth rate of K562 cells remained unchanged or even accelerated according to CCK-8 absorbance measurements (Fig. 6C). Moreover, microscopy observations confirmed the presence of heterotypic leukemia-stroma spheroids visualized by the fluorescently tracked cells (HS-5-cyan; K562-magenta), indicating good co-culture conditions even upon imatinib treatment (Supplementary Figure S2).

Altogether, these data confirm the significant stroma-mediated protection from imatinib and demonstrate that the proposed in vitro 3D model can be utilized for drug screening under leukemic microenvironment conditions.

## Discussion

In summary, we propose a microfluidic method of reproducible high-throughput formulation of leukemia-stroma-ECM microtissues that recapitulate some of the critical features of the chronic myeloid leukemia microenvironment. We use droplet microfluidics to co-encapsulate the leukemic and bone-marrow stromal cells inside GelMA hydrogel microbeads at a high rate (around 12 000/min) and used the generated leukemic microtissues to test the impact of stroma on the development of drug resistance.

Our study shows that GelMA provides the best conditions for co-culture and spheroid formation. In particular, GelMA microgels have been previously employed as highly reproducible 3D culture vehicles, and used, e.g., in skin, bone, and liver microtissue engineering (Tan et al., 2021; Wang et al., 2019; Zhao et al., 2016b) as well as in drug screening using prostate-and breast cancer microtissues which included investigation of the effect of 3D culture on drug resistance (Antunes et al., 2019; Yang et al., 2020). Formation of microtissues inside GelMA microbeads has been demonstrated previously, e.g., with the use of colon cancer cells and fibroblasts (Mohamed et al., 2019; Zhang et al., 2022), yet only in the case of monocultures. Here, we develop a more advanced model of leukemic cancer microenvironment, including also the stromal cells. We establish that 4.8% GelMA hydrogel provides optimal conditions for both cancer and stromal cell proliferation, whereas the addition of other biopolymers such as HAMA or fibrin to GelMA appears disadvantageous either due to slower proliferation (GelMA+HAMA) or poor crosslinking (GelMA+fibrin), in the latter case associated with elevated cytotoxicity and/or cell outmigration. With precisely established conditions, we propose a reproducible 3D leukemic CME model including all of its basic components, that is the leukemic cells, the bone marrow cells, and the extracellular matrix. We observe the formation of co-culture leukemic-stroma spheroids inside the microbeads within 48h post-encapsulation and find that the generated cancer microtissues develop stroma-mediated protection from Imatinib, which confirms the expected effect of the stromal barrier on drug sensitivity.

Non-high-throughput leukemic microenvironment models have been proposed previously. 3D tissue-engineered bone marrow closely recapitulated pathophysiological conditions in multiple myeloma and has been proposed as a feasible platform to predict clinical drug efficacy (Alhallak et al., 2021). The patient-derived 3D models of the bone marrow microenvironment of multiple myeloma for personalized medicine were recently rediscussed with all limitations and remaining challenges (Lourenço et al., 2022). The potential of 3D in vitro models has been also underlined in the context of understanding drug resistance in leukemia stem cells (Al-Kaabneh et al., 2022).

In fact, stroma-mediated protection seems to be one of the most important current clinical challenges in leukemia therapy. Therefore, the implementation of adequate models mimicking the bone marrow niche is critical for successful drug development and testing. Despite the superior physiological relevance of the 3D culture, reliable 3D leukemia CME models remain scarce. Recently, James et al. (James et al., 2023) reported a 3D cell culture model with the leukemic cells embedded in an external peptide-based hydrogel mimicking the properties of the ECM. Authors used multi-well plates to study the responses of various types of leukemic cell lines embedded in a peptide hydrogel layer to several anti-cancer drugs supplied from the external media. The composition of the hydrogel was optimized in terms of the efficiency of the formation of the leukemic cellular colonies; however, no stromal cells were directly applied in the model. Also, the dispersion of cells in a bulk hydrogel, despite the ease of application, had its limitation in terms of further manipulation of the individual ‘colony forming units’, i.e., the cancer microtissues. In general, the macro-encapsulation approaches such as those proposed by James et al. or others (Belloni et al., 2022) have a drawback in that they do not allow aspiration of the individual microtissues, nor their further transfer or plating. Also, the precision confocal imaging remains limited due to the large thickness of the hydrogel samples (typically ∼1 mm) and the diffraction of light by the surrounding co-embedded cells and/or the hydrogel itself. Recently, an advanced bone marrow microenvironment model—yet, non-leukemic— utilizing organ-on-chip technology has been proposed (Sharipol et al., 2022). However, the approach was strongly limited in terms of throughput (one tissue per chip) which provides an obstacle in drug screening studies requiring very high throughput.

Here, we demonstrate an automated and highly reproducible micro-encapsulation approach relying on the dispersion of hydrogel-cell mixture into thousands of monodisperse hydrogel microbeads. Reproducible formation of the independent leukemic microenvironments encapsulated in separate microbeads allows for efficient diffusion of nutrients and drugs from the media and precise modeling of cellular responses including the development of drug resistance. To assess the performance of the model in comparison with the previous macro-encapsulation approaches mentioned above, we have performed an auxiliary experiment employing the conventional 3D matrigel leukemia-bone marrow spheroid model (bulk 1:1 co-culture). We have found that such a bulk model is not capable of reproducing the physiological stroma-mediated resistance to Imatinib (Supplementary Fig. S3).

We propose that our droplet-based 3D co-culture model of the leukemic microenvironment described here could be further developed for more advanced biological systems. First, we note that microbeads could be also grown under hypoxia. Therefore, our model could be used to develop an efficient therapy against leukemic stem-and progenitor cells which settle in the bone marrow and are highly resistant to apoptosis. Also, the model might be easily extended to incorporate other cell types, e.g., to mimic different specialized lymphoid tissues and/or interactions with immune cells. Those could include 3D models based on the co-culture of organoids and immune cells, thereby presenting a novel approach for tumor immunology study, as recently demonstrated by Mu et al. (Mu et al., 2023). The more complex 3D tri-culture model of leukemia, including mesenchymal stem/stromal and HSPC cells mimicking the leukemic bone marrow, yet without ECM, has been recently presented as a platform for drug testing and discovery of novel anti-leukemic therapy (Zippel et al., 2022). Our microbead-based leukemia bone marrow microenvironment model, incorporating the ECM and amenable to high-throughput processing, could therefore pave the way for more complex, heterotypic cancer microenvironment models with prospective translational and clinical potential.

In fact, till now, only the *in vivo*/*ex vivo* tests allowed for studies of leukemic bone marrow niche under physiological conditions (Lourenco et al., 2022). In line with the recent FDA regulations which approved drug development without animal studies (Adashi et al., 2023), the future drug development strategies are supposed to limit the *in vivo* models. This new direction in preclinical research strongly encourages the development of highly efficient 3D physiological models.

In summary, the microfluidic method of formulation of the leukemic microtissues that we propose in this work warrants high throughput of production, low consumption of reagents, and excellent reproducibility, and thus lays the ground for the development of future drug screening platforms for rapid evaluation of anti-leukemia therapeutics. Our leukemic microtissues fill the gap between preclinical studies and animal models and, upon further development—e.g., trapping of cancer microtissues inside a microfluidic chip for long-term culture under controlled perfusion (Bouquerel et al., 2023)—could provide for a highly efficient drug testing platform in leukemia.

## Materials and methods

### Cell culture

The human chronic myeloid leukemia K562 (#CCL-243) cells and human stromal cell line HS-5 (#CRL-11882) were purchased from American Type Culture Collection. The cells were cultured in RPMI 1640, supplemented with 10 % fetal bovine serum and 1 % mixture of antibiotics penicillin-streptomycin (all from Gibco, Waltham, MA, USA) at 5 % CO_2_ and 37 °C in a humidified chamber. K562 cells were grown in suspension culture and cells were seeded at a concentration of 300,000 viable cells/ml. The culture was either re-fed with fresh medium every three days or split again. HS-5 cells were grown to confluence and harvested by trypsinization, using a 0.25 mg/mL trypsin/EDTA (ThermoFisher, Waltham, MA, USA) and resuspended in the fresh culture medium. Following Trypan Blue staining, K562 and HS-5 cell lines were enumerated on the Countess II FL Automated Cell Counter (ThermoFisher, Waltham, MA, USA). The cell lines were tested for mycoplasma contamination using a PCR-based approach.

### GelMA, HAMA synthesis and hydrogel preparation

Gelatin methacryloyl (GelMA), a photo-crosslinkable hydrogel based on gelatin, was prepared using a previously described protocol (Yue et al., 2015). Firstly, 10 g gelatin, type A, Bloom value = 300 (Sigma Aldrich, Saint Louis, MO, USA) was dissolved in 100 ml phosphate-buffered saline (PBS; buffer pH = 7.4; Sigma-Aldrich, USA) at 50°C with continuous stirring. After having the gelatin completely dissolved, methacrylic anhydride (MA; Sigma-Aldrich, USA) was added to the mixture (0.8 ml per 1 g of gelatin) and stirred for 5-6 h, 50°C. Next, the solution was diluted with warm PBS (40°C) and dialyzed (with 12-kDa molecular weight cut-off membrane; Sigma Aldrich, USA) against distilled water for 5-7 days at RT to remove excess methacrylic acid and other impurities. Finally, the solution was freeze-dried and stored at −°C. The freeze-dried GelMA was used to form 4% and 6% (w/v) solution in 1X PBS by stirring for 1h at 50°C.

For the GelMA-HAMA hydrogel, 1% solution of methacrylated hyaluronic acid (HAMA) (Polbionica sp. Zoo., Poland) was prepared in 1x PBS. The mixture was stirred (1000 rpm) at 4°C until the methacrylate dissolved. The HAMA solution was mixed in 1:1 ratio with 12% GelMA solution in PBS to prepare 6% GelMA with 0.5% HAMA solution.

Additionally, 5 mg/mL of fibrinogen (Sigma-Aldrich, St. Louis, MO, USA) stock was prepared and mixed in a 1:1 ratio with 12 % GelMA (v/v) to obtain a final solution of 0.25 % fibrin (v/v) in 6% GelMA (v/v). Similarly, a 0.1 % concentration of fibrin in 6 % GelMA was obtained by mixing the stock in a 1:4 ratio with 7.5 % GelMA. The fibrinogen was converted to fibrin with 0.625 units of thrombin (Sigma-Aldrich, St. Louis, MO, USA) added in a volume fraction of 9:1 (fibrinogen to thrombin) prior to hydrogel photocrosslinking.

Hydrogel samples (3 replicates of each condition) were obtained by exposing 100 µl of an aqueous solution of GelMA, GelMA-HAMA or GelMA-fibrin prepolymers with added photoinitiator (0.4 % v/v) Lithium phenyl-2,4,6-trimethylbenzoylphosphinate (LAP) (Cellink, Gothenburg, Sweden) to visible violet light (405 nm, 10 sec exposition) using a 96-well plate and a custom 405 nm LED array (5 x 5 = 25 LEDs at 5 mm spacing, each with the output power P = 1W). The irradiance was 280 mW/cm^2^ as measured by PM 100D (Thorlabs, Inc., United States) optical power meter placed 1 cm above the center of the LED array. The lamp was placed in the PMMA box with a sliding door fitted with a UV-protectant screen.

The cells (K562 and HS-5 in monoculture) were intermixed with hydrogel precursor prior to crosslinking. For the intermixing, a cell suspension of cells with culture media was mixed in a 1:4 ratio with the different hydrogel compositions. The resulting final hydrogel compositions were accordingly as follows: 3.2% GelMA, 4.8% GelMA, 4.8% GelMA with 0.4% HAMA, 4.8% GelMA with 0.08% Fibrin and 4.8% GelMA with 0.2% Fibrin.

### Cell Counting Kit-8 (CCK-8) assay

The colorimetric Cell Counting Kit-8 (CCK-8; MedchemExpress, NJ, USA) assay assessed cell proliferation at 24, 48, and 72 h of treatment. After cell encapsulation, microbeads were seeded onto the 96-well plate in 4 replicates. Encapsulated cells were cultured for 24h and then the medium was replaced with a fresh medium with 0.5-2 µM imatinib every day for 1-3 days. 10 μL of CCK-8 solution was added to the 100 μL medium in each well and incubated at 37°C for 4 h. Viable cells were identified relying on the ability of mitochondrial dehydrogenases to oxidize water-soluble tetrazolium 8 (WST-8) into a formazan product. The absorbance of the sample at 450 nm was measured with a Synergy HTX Microplate Spectrophotometer (BioTek, Winooski, VT, USA), and plotted to obtain cell growth curves. The well with medium without microbeads was used as the negative control.

### Cell Tracker (red/green fluorescence) labeling

The cells were harvested, washed with Dulbecco’s phosphate-buffered saline (DPBS; ThermoFisher, Waltham, MA, USA), and resuspended in serum-free medium with the cell tracking dyes. K562 cells were labeled with CellTracker™ Red CMTPX Dye (Ex/Em 577/602 nm) with 1 μL from the 10 mM dye stock added to 2 mL of serum-free media. Similarly, CellTracker™ Green CMFDA Dye (Ex/Em 492/ 517 nm; Thermo Fisher, Waltham, MA, USA) was used to label the HS-5 cells. The cells were incubated at 37°C with 5% CO_2_ in a humidified chamber for 30 min with tracking dye. Next, the staining solution was removed, and complete media was added. Cells were counted using Countess II FL Automated Cell Counter (ThermoFisher, Waltham, MA, USA) and the 3-6 x 10^7^/ml cell stock was prepared for further experiments. The dyes allowed visualization of both cell types with different colors under fluorescence microscopy. CellTracker probes freely pass through cell membranes into the cells where they get transformed into a cell-impermeant, fluorescent product –retained in living cells through several generations.

### LIVE/DEAD staining

A LIVE/DEAD™ Viability/Cytotoxicity Kit for mammalian cells (Thermo Fisher, USA) was used to stain the live and dead cells. The Calcein-AM component (Ex/Em = 495/515 nm) stained viable cells green by permeating the cell membrane and enzymatically degrading to a fluorescent green calcein product, while the BOBO-3 Iodide component (Ex/Em = 528/617 nm) permeated only damaged cell membranes and stained them red when bound to nucleic DNA.

The media were aspirated by gentle suction and replaced with DPBS containing LIVE/DEAD dyes (1:5), then incubated in a cell culture incubator for 15 min. The microscopy images were acquired using the Nikon Eclipse TSR2 (Nikon, Tokyo, Japan) microscope (typically using 10× magnification objective) and analyzed with Fiji (ImageJ).

### Cell Proliferation Dye eFluor450 labeling for flow cytometry analyses

For flow cytometry experiments, Cell Proliferation Dye eFluor™450 (Invitrogen) was used to label HS-5 cells before setting up a co-culture with K562 cells. Cells were trypsinized and washed twice with PBS. The cell pellet was resuspended in 1 ml of 10µM Cell Proliferation Dye eFluor™450 solution (prepared in PBS) and incubated for 10 min at 37°C in the dark. Labeling was stopped by adding cold complete media and was followed by 5 min incubation on ice. Then, cells were washed twice with complete media, counted, and prepared for cell encapsulation.

### Apoptosis analysis by flow cytometry

Apoptosis was determined by flow cytometry using the Annexin V-FITC/7-AAD Apoptosis Detection Kit (BD Biosciences) according to the manufacturer’s instructions. The percentage of Annexin-V-positive cells was calculated as a percentage of apoptotic cells. K562 and HS-5 cells were separated by gating based on Proliferation Dye eFluor450 fluorescence (HS-5 eFluor450-positive and K562 eFluor450-negative populations).

### Mitochondrial mass analysis by flow cytometry

Mitochondrial mass was analyzed by flow cytometry using the MitoTracker™ Green FM Dye (Invitrogen). 0.5 x 10^6^ cells were transferred to the FACS tubes. Cells were washed twice with PBS, then resuspended in 100µL of 200nM MitoTracker™ Green FM Dye solution (prepared in serum-free media). Then, cells were incubated for 30 minutes at 37°C. After that, cells were washed twice with PBS and analyzed with BD LSR Fortessa cytometer (BD Biosciences). K562 and HS-5 cells were separated by gating based on Proliferation Dye eFluor450 fluorescence (HS-5 eFluor450-positive and K562 eFluor450-negative populations).

### Microfluidic chip fabrication

The microchip was designed using AutoCAD software and channels were micro-milled in polymethylmethacrylate (PMMA; b/b ex 5mm – clear, extruded, Rokad Sp. Z o.o., Białebłoto-Stara Wieś, Długosiodło, Poland). Polydimethylsiloxane (PDMS) (SYLGARD™ 184 Silicone Elastomer Kit, Dow Corning Corporation, Michigan, United States) was used to fabricate the copy of microfluidic chips. A 1:9 w/w ratio of the curing agent and PDMS pre-polymer was mixed, degassed under vacuum, and poured onto the molds. PDMS was cured for 12 h at 80 °C. PDMS replicates from molds were exposed to oxygen plasma (29.6 W High RF Power) for 2 minutes in a vacuum-pressurized (<300 mTorr) plasma cleaner device (PDC-002-CE; Harrick Plasma, NY, USA) and silanized with trichloro(1H,1H,2H,2H-perfluorooctyl)silane (PFOCTS; Sigma-Aldrich, Saint Louis, Missouri, United States). The geometry of the microchannels used for droplet generation and optimization of the applied flow rates will be described in detail elsewhere. Shortly, the chip consisted of an inlet for the droplet phase supplying multiple droplet generators operating via so-called step-emulsification (Sugiura et al., 2001), and an outlet for the generated emulsion.

### Cell encapsulation and drug treatment

4-12 x 10^6^ K652 or HS-5 (K625: HS-5 1:1 for co-culture experiments) cells (resuspended in 200 µl culture medium) were mixed with 800 µl hydrogel (cell-mixture: hydrogel ratio 1:4) and dispersed into droplets using the microfluidic device supplied with fluorinated oil as the external phase. For that purpose, we used Novec™ 7500 (3M, USA) with added 2 % (w/w) fluorosurfactant (PFPE-PEG-PFPE; Chemipan R&D Laboratories, Warsaw, Poland). The flows of the cell-hydrogel and the oil phases were controlled by a double-channel syringe pump (33 DDS Pump, Harvard Apparatus, Holliston, Massachusetts, USA) and supplied with 1 ml and 5 ml plastic BD syringes, respectively. The generated water-in-oil (W/O) emulsion was collected in a 40 mm diameter Petri dish and placed directly at the custom-built 405 nm LED panel. The LED panel was provided by the Foundation for Research and Science Development (FBiRN, Warsaw, Poland) and consisted of a 5 x 5 array of 405 nm LEDs, each with the characteristic radiant flux Φ_e_ = 1100 mW. We illuminated the emulsion for 10 sec at the irradiance of 280 mW/cm^2^. The crosslinked microbeads suspended in oil were subsequently transferred into eppendorf tubes and the oil from the bottom was aspirated and discarded. Next, 500 µl of Novec 7500 containing 20 % 1H,1H,2H,2H-perfluorooctanol (PFO; ThermoFisher Scientific GmBH, Germany) was added to the suspension and mixed by pipetting resulting in bead clustering. This is a standard procedure (Mao et al., 2017) wherein PFO removes the surfactant from the surface of the beads. Novec 7500 with PFO was subsequently aspirated and discarded and replaced with pure Novec 7500. This washing step was repeated three times. Finally, Novec 7500 was removed, replaced with 1 ml PBS, and centrifuged at 1100 rpm for 5 minutes. After centrifugation three distinct layers (oil, microbeads, and PBS, counting from the bottom) could be distinguished. The oil layer and the PBS layer were subsequently removed leaving only the clustered beads in the eppendorf tube. This PBS washing process was repeated two times to remove the remaining oil. After the final wash, PBS was replaced with fresh cell culture media (using the same volume for each sample), transferred to 96-well cell culture plates, and cultured at 5 % CO_2_ and 37°C in a humidified incubator.

The encapsulated cells were cultured for 24h after which the medium was exchanged to a fresh medium containing 0.5-2.0 µM imatinib (kindly provided by the Lukasiewicz Research Network – Industrial Chemistry Institute, formerly Pharmaceutical Institute, Warsaw, Poland). The cells were subsequently cultured for another 24-72 h and observed under the Nikon Eclipse TSR2 fluorescence microscope or Nikon A1R confocal microscope (Nikon, Tokyo, Japan).

### Cell releasing from microbeads

The microbeads were treated with 1mg/ml of collagenase (Thermo Fisher, Waltham, Massachusetts, USA) for 7 min at 37°C in a humidified incubator until beads were dissolved. The process was controlled under a Nikon fluorescence microscope using 10x magnification and the cell condition was verified using LIVE/DEAD staining.

### Statistical Analysis

All experiments were repeated at least three times. Data are reported as mean ± SD. Data were analyzed using one-way ANOVA, followed by Dunnet’s or Mann-Whitney tests (GraphPad, Prism 6.00 for Windows, Graf Pad Software, San Diego, CA, USA). The *p*-value of <0.05 was considered statistically significant with *, *p* < 0.01 with **, *p* < 0.001 with ***, and *p* < 0.0001 with ****.

## Author contributions

MRR developed methods, designed and performed experiments, analyzed data, prepared graphs and prepared draft of the manuscript. DM performed experiments, analyzed data, and was involved in graphs and manuscript preparation. LT-K was involved in experiment design, performing, and data analysis as well as manuscript preparation. PP-B performed in vitro 3D matrigel experiments, analyzed the data, and revised the manuscript. KP and JG conceptualized the project, designed and supervised experiments and data analysis, prepared the draft of the manuscript. All authors have read, edited, and agreed to the final version of the manuscript.

## Acknowledgements

This work was supported by National Science Centre grants: 2021/41/B/NZ5/04077 (KP), PRELUDIUM BIS 2019/35/O/NZ3/01936 (KP, LT-K), OPUS 2019/33/B/ST8/02145 (DM, JG).

## References

1. Adamo, A., Delfino, P., Gatti, A., Bonato, A., Takam Kamga, P., Bazzoni, R., Ugel, S., Mercuri, A., Caligola, S., Krampera, M., 2020. HS-5 and HS-27A Stromal Cell Lines to Study Bone Marrow Mesenchymal Stromal Cell-Mediated Support to Cancer Development. Front. Cell Dev. Biol. 8, 584232. 10.3389/fcell.2020.584232

2. Adashi, E.Y., O’Mahony, D.P., Cohen, I.G., 2023. The FDA Modernization Act 2.0: Drug Testing in Animals is Rendered Optional. The American Journal of Medicine 136, 853–854. 10.1016/j.amjmed.2023.03.033

3. Agarwal, P., Zhang, B., Ho, Y., Cook, A., Li, L., Mikhail, F.M., Wang, Y., McLaughlin, M.E., Bhatia, R., 2017. Enhanced targeting of CML stem and progenitor cells by inhibition of porcupine acyltransferase in combination with TKI. Blood 129, 1008–1020. 10.1182/blood-2016-05-714089

4. Alhallak, K., Jeske, A., De La Puente, P., Sun, J., Fiala, M., Azab, F., Muz, B., Sahin, I., Vij, R., DiPersio, J.F., Azab, A.K., 2021. A pilot study of 3D tissue-engineered bone marrow culture as a tool to predict patient response to therapy in multiple myeloma. Sci Rep 11, 19343. 10.1038/s41598-021-98760-9

5. Al-Kaabneh, B., Frisch, B., Aljitawi, O.S., 2022. The Potential Role of 3D In Vitro Acute Myeloid Leukemia Culture Models in Understanding Drug Resistance in Leukemia Stem Cells. Cancers 14, 5252. 10.3390/cancers14215252

6. Antunes, J., Gaspar, V.M., Ferreira, L., Monteiro, M., Henrique, R., Jerónimo, C., Mano, J.F., 2019. In-air production of 3D co-culture tumor spheroid hydrogels for expedited drug screening. Acta Biomaterialia 94, 392–409. 10.1016/j.actbio.2019.06.012

7. Aoyagi, M., Furusawa, S., Waga, K., Tsunogake, S., Shishido, H., 1994. Suppression of Normal Hematopoiesis in Acute Leukemia: Effect of Leukemic Cells on Bone Marrow Stromal Cells and Hematopoietic Progenitor Cells. Intern. Med. 33, 288–295. 10.2169/internalmedicine.33.288

8. Araujo-Ayala, F., Dobaño-López, C., Valero, J.G., Nadeu, F., Gava, F., Faria, C., Norlund, M., Morin, R., Bernes-Lasserre, P., Serrat, N., Playa-Albinyana, H., Giménez, R., Campo, E., Lagarde, J.-M., López-Guillermo, A., Gine, E., Colomer, D., Bezombes, C., Pérez-Galán, P., 2023. A novel patient-derived 3D model recapitulates mantle cell lymphoma lymph node signaling, immune profile and in vivo ibrutinib responses. Leukemia 37, 1311–1323. 10.1038/s41375-023-01885-1

9. Belloni, D., Ferrarini, M., Ferrero, E., Guzzeloni, V., Barbaglio, F., Ghia, P., Scielzo, C., 2022. Protocol for generation of 3D bone marrow surrogate microenvironments in a rotary cell culture system. STAR Protocols 3, 101601. 10.1016/j.xpro.2022.101601

10. Bhamidipati, P.K., Kantarjian, H., Cortes, J., Cornelison, A.M., Jabbour, E., 2013. Management of imatinib-resistant patients with chronic myeloid leukemia. Therapeutic Advances in Hematology 4, 103–117. 10.1177/2040620712468289

11. Bouquerel, C., Dubrova, A., Hofer, I., Phan, D.T.T., Bernheim, M., Ladaigue, S., Cavaniol, C., Maddalo, D., Cabel, L., Mechta-Grigoriou, F., Wilhelm, C., Zalcman, G., Parrini, M.C., Descroix, S., 2023. Bridging the gap between tumor-on-chip and clinics: a systematic review of 15 years of studies. Lab Chip 23, 3906–3935. 10.1039/D3LC00531C

12. Camci-Unal, G., Cuttica, D., Annabi, N., Demarchi, D., Khademhosseini, A., 2013. Synthesis and Characterization of Hybrid Hyaluronic Acid-Gelatin Hydrogels. Biomacromolecules 14, 1085– 1092. 10.1021/bm3019856

13. Chen, Z., Zheng, Y., Yang, Y., Kang, J., You, M.J., Tian, C., 2021. Abnormal bone marrow microenvironment: the “harbor” of acute lymphoblastic leukemia cells. Blood Science 3, 29–34. 10.1097/BS9.0000000000000071

14. Civini, S., Jin, P., Ren, J., Sabatino, M., Castiello, L., Jin, J., Wang, H., Zhao, Y., Marincola, F., Stroncek, D., 2013. Leukemia cells induce changes in human bone marrow stromal cells. J Transl Med 11, 298. 10.1186/1479-5876-11-298

15. Colmone, A., Amorim, M., Pontier, A.L., Wang, S., Jablonski, E., Sipkins, D.A., 2008. Leukemic Cells Create Bone Marrow Niches That Disrupt the Behavior of Normal Hematopoietic Progenitor Cells. Science 322, 1861–1865. 10.1126/science.1164390

16. De Rutte, J.M., Koh, J., Di Carlo, D., 2019. Scalable High-Throughput Production of Modular Microgels for In Situ Assembly of Microporous Tissue Scaffolds. Adv Funct Materials 29, 1900071. 10.1002/adfm.201900071

17. Deininger, M.W.N., Goldman, J.M., Lydon, N., Melo, J.V., 1997. The Tyrosine Kinase Inhibitor CGP57148B Selectively Inhibits the Growth of BCR-ABL–Positive Cells. Blood 90, 3691–3698. 10.1182/blood.V90.9.3691

18. Dolinska, M., Cai, H., Mansson, A., Shen, J., Xiao, P., Bouderlique, T., Li, X., Leonard, E., Chang, M., Gao, Y., Medina Giménez, J.P., Kondo, M., Sandhow, L., Johansson, A.-S., Deneberg, S., Söderlund, S., Jädersten, M., Ungerstedt, J.S., Tobiasson, M., Östman, A., Mustjoki, S., Stenke, L., Le Blanc, K., Hellstrom-Lindberg, E.S., Lehmann, S., Ekblom, M., Olsson-Strömberg, U., Sigvardsson, M., Qian, H., 2023. Characterization of Bone Marrow Niche in Chronic Myeloid Leukemia Patients Identifies CXCL14 as a New Therapeutic Option. Blood blood.2022016896. 10.1182/blood.2022016896

19. Druker, B.J., Talpaz, M., Resta, D.J., Peng, B., Buchdunger, E., Ford, J.M., Lydon, N.B., Kantarjian, H., Capdeville, R., Ohno-Jones, S., Sawyers, C.L., 2001. Efficacy and Safety of a Specific Inhibitor of the BCR-ABL Tyrosine Kinase in Chronic Myeloid Leukemia. N Engl J Med 344, 1031–1037. 10.1056/NEJM200104053441401

20. Druker, B.J., Tamura, S., Buchdunger, E., Ohno, S., Segal, G.M., Fanning, S., Zimmermann, J., Lydon, N.B., 1996. Effects of a selective inhibitor of the Abl tyrosine kinase on the growth of Bcr–Abl positive cells. Nat Med 2, 561–566. 10.1038/nm0596-561

21. Duan, C.-W., Shi, J., Chen, J., Wang, B., Yu, Y.-H., Qin, X., Zhou, X.-C., Cai, Y.-J., Li, Z.-Q., Zhang, F., Yin, M.-Z., Tao, Y., Mi, J.-Q., Li, L.-H., Enver, T., Chen, G.-Q., Hong, D.-L., 2014. Leukemia Propagating Cells Rebuild an Evolving Niche in Response to Therapy. Cancer Cell 25, 778–793. 10.1016/j.ccr.2014.04.015

22. Federica Barbaglio, Daniela Belloni, Lydia Scarfò, Francesca Vittoria Sbrana, Maurilio Ponzoni, Lucia Bongiovanni, Luca Pavesi, Desiree Zambroni, Kostas Stamatopoulos, Valeria R. Caiolfa, Elisabetta Ferrero, Paolo Ghia, Cristina Scielzo, 2020. Three-dimensional co-culture model of chronic lymphocytic leukemia bone marrow microenvironment predicts patient-specific response to mobilizing agents. haematol 106, 2334–2344. 10.3324/haematol.2020.248112

23. Fontana, F., Marzagalli, M., Sommariva, M., Gagliano, N., Limonta, P., 2021. In Vitro 3D Cultures to Model the Tumor Microenvironment. Cancers 13, 2970. 10.3390/cancers13122970

24. Germain, N., Dhayer, M., Dekiouk, S., Marchetti, P., 2022. Current Advances in 3D Bioprinting for Cancer Modeling and Personalized Medicine. IJMS 23, 3432. 10.3390/ijms23073432

25. Habanjar, O., Diab-Assaf, M., Caldefie-Chezet, F., Delort, L., 2021. 3D Cell Culture Systems: Tumor Application, Advantages, and Disadvantages. IJMS 22, 12200. 10.3390/ijms222212200

26. Hantschel, O., Grebien, F., Superti-Furga, G., 2012. The Growing Arsenal of ATP-Competitive and Allosteric Inhibitors of BCR–ABL. Cancer Research 72, 4890–4895. 10.1158/0008-5472.CAN-12-1276

27. Hermansen, J.U., Yin, Y., Urban, A., Myklebust, C.V., Karlsen, L., Melvold, K., Tveita, A.A., Taskén, K., Munthe, L.A., Tjønnfjord, G.E., Skånland, S.S., 2023. A tumor microenvironment model of chronic lymphocytic leukemia enables drug sensitivity testing to guide precision medicine. Cell Death Discov. 9, 125. 10.1038/s41420-023-01426-w

28. Im, G.-B., Lin, R.-Z., 2022. Bioengineering for vascularization: Trends and directions of photocrosslinkable gelatin methacrylate hydrogels. Front. Bioeng. Biotechnol. 10, 1053491. 10.3389/fbioe.2022.1053491

29. Jabbour, E., Parikh, S.A., Kantarjian, H., Cortes, J., 2011. Chronic Myeloid Leukemia: Mechanisms of Resistance and Treatment. Hematology/Oncology Clinics of North America 25, 981–995. 10.1016/j.hoc.2011.09.004

30. Jabbour, E.J., Cortes, J.E., Kantarjian, H.M., 2013. Resistance to Tyrosine Kinase Inhibition Therapy for Chronic Myelogenous Leukemia: A Clinical Perspective and Emerging Treatment Options. Clinical Lymphoma Myeloma and Leukemia 13, 515–529. 10.1016/j.clml.2013.03.018

31. James, J.R., Curd, J., Ashworth, J.C., Abuhantash, M., Grundy, M., Seedhouse, C.H., Arkill, K.P., Wright, A.J., Merry, C.L.R., Thompson, A., 2023. Hydrogel-Based Pre-Clinical Evaluation of Repurposed FDA-Approved Drugs for AML. IJMS 24, 4235. 10.3390/ijms24044235

32. Joshi, S.K., Nechiporuk, T., Bottomly, D., Piehowski, P.D., Reisz, J.A., Pittsenbarger, J., Kaempf, A., Gosline, S.J.C., Wang, Y.-T., Hansen, J.R., Gritsenko, M.A., Hutchinson, C., Weitz, K.K., Moon, J., Cendali, F., Fillmore, T.L., Tsai, C.-F., Schepmoes, A.A., Shi, T., Arshad, O.A., McDermott, J.E., Babur, O., Watanabe-Smith, K., Demir, E., D’Alessandro, A., Liu, T., Tognon, C.E., Tyner, J.W., McWeeney, S.K., Rodland, K.D., Druker, B.J., Traer, E., 2021. The AML microenvironment catalyzes a stepwise evolution to gilteritinib resistance. Cancer Cell 39, 999–1014.e8. 10.1016/j.ccell.2021.06.003

33. Klotz, B.J., Gawlitta, D., Rosenberg, A.J.W.P., Malda, J., Melchels, F.P.W., 2016. Gelatin-Methacryloyl Hydrogels: Towards Biofabrication-Based Tissue Repair. Trends in Biotechnology 34, 394–407. 10.1016/j.tibtech.2016.01.002

34. Kolba, M.D., Dudka, W., Zaręba-Kozioł, M., Kominek, A., Ronchi, P., Turos, L., Chroscicki, P., Wlodarczyk, J., Schwab, Y., Klejman, A., Cysewski, D., Srpan, K., Davis, D.M., Piwocka, K., 2019. Tunneling nanotube-mediated intercellular vesicle and protein transfer in the stroma-provided imatinib resistance in chronic myeloid leukemia cells. Cell Death Dis 10, 817. 10.1038/s41419-019-2045-8

35. Korn, C., Méndez-Ferrer, S., 2017. Myeloid malignancies and the microenvironment. Blood 129, 811– 822. 10.1182/blood-2016-09-670224

36. Kumar, A., Bhattacharyya, J., Jaganathan, B.G., 2017. Adhesion to stromal cells mediates imatinib resistance in chronic myeloid leukemia through ERK and BMP signaling pathways. Sci Rep 7, 9535. 10.1038/s41598-017-10373-3

37. Le, B.V., Podszywalow-Bartnicka, P., Maifrede, S., Sullivan-Reed, K., Nieborowska-Skorska, M., Golovine, K., Yao, J.-C., Nejati, R., Cai, K.Q., Caruso, L.B., Swatler, J., Dabrowski, M., Lian, Z., Valent, P., Paietta, E.M., Levine, R.L., Fernandez, H.F., Tallman, M.S., Litzow, M.R., Huang, J., Challen, G.A., Link, D., Tempera, I., Wasik, M.A., Piwocka, K., Skorski, T., 2020. TGFβR-SMAD3 Signaling Induces Resistance to PARP Inhibitors in the Bone Marrow Microenvironment. Cell Reports 33, 108221. 10.1016/j.celrep.2020.108221

38. Liu, J., Tan, Y., Zhang, H., Zhang, Y., Xu, P., Chen, J., Poh, Y.-C., Tang, K., Wang, N., Huang, B., 2012. Soft fibrin gels promote selection and growth of tumorigenic cells. Nature Mater 11, 734– 741. 10.1038/nmat3361

39. Lourenço, D., Lopes, R., Pestana, C., Queirós, A.C., João, C., Carneiro, E.A., 2022. Patient-Derived Multiple Myeloma 3D Models for Personalized Medicine—Are We There Yet? IJMS 23, 12888. 10.3390/ijms232112888

40. Mao, A.S., Shin, J.-W., Utech, S., Wang, H., Uzun, O., Li, W., Cooper, M., Hu, Y., Zhang, L., Weitz, D.A., Mooney, D.J., 2017. Deterministic encapsulation of single cells in thin tunable microgels for niche modelling and therapeutic delivery. Nature Mater 16, 236–243. 10.1038/nmat4781

41. Melo, J.V., Chuah, C., 2007. Resistance to imatinib mesylate in chronic myeloid leukaemia. Cancer Letters 249, 121–132. 10.1016/j.canlet.2006.07.010

42. Mohamed, M.G.A., Kheiri, S., Islam, S., Kumar, H., Yang, A., Kim, K., 2019. An integrated microfluidic flow-focusing platform for on-chip fabrication and filtration of cell-laden microgels. Lab Chip 19, 1621–1632. 10.1039/C9LC00073A

43. Moutinho, S., 2023. Researchers and regulators plan for a future without lab animals. Nat Med s41591–023-02362-z. 10.1038/s41591-023-02362-z

44. Mu, P., Zhou, S., Lv, T., Xia, F., Shen, L., Wan, J., Wang, Y., Zhang, H., Cai, S., Peng, J., Hua, G., Zhang, Z., 2023. Newly developed 3D in vitro models to study tumor–immune interaction. J Exp Clin Cancer Res 42, 81. 10.1186/s13046-023-02653-w

45. Nichol, J.W., Koshy, S.T., Bae, H., Hwang, C.M., Yamanlar, S., Khademhosseini, A., 2010. Cell-laden microengineered gelatin methacrylate hydrogels. Biomaterials 31, 5536–5544. 10.1016/j.biomaterials.2010.03.064

46. Pal, D., Blair, H., Parker, J., Hockney, S., Beckett, M., Singh, M., Tirtakusuma, R., Nelson, R., McNeill, H., Angel, S.H., Wilson, A., Nizami, S., Nakjang, S., Zhou, P., Schwab, C., Sinclair, P., Russell, L.J., Coxhead, J., Halsey, C., Allan, J.M., Harrison, C.J., Moorman, A.V., Heidenreich, O., Vormoor, J., 2022. hiPSC-derived bone marrow milieu identifies a clinically actionable driver of niche-mediated treatment resistance in leukemia. Cell Reports Medicine 3, 100717. 10.1016/j.xcrm.2022.100717

48. Park, H.J., Gregory, M.A., Zaberezhnyy, V., Goodspeed, A., Jordan, C.T., Kieft, J.S., DeGregori, J., 2022. Therapeutic resistance in acute myeloid leukemia cells is mediated by a novel ATM/mTOR pathway regulating oxidative phosphorylation. eLife 11, e79940. 10.7554/eLife.79940

49. Patterson, S.D., Copland, M., 2023. The Bone Marrow Immune Microenvironment in CML: Treatment Responses, Treatment-Free Remission, and Therapeutic Vulnerabilities. Curr Hematol Malig Rep 18, 19–32. 10.1007/s11899-023-00688-6

50. Podszywalow-Bartnicka, P., Cmoch, A., Wolczyk, M., Bugajski, L., Tkaczyk, M., Dadlez, M., Nieborowska-Skorska, M., Koromilas, A.E., Skorski, T., Piwocka, K., 2016. Increased phosphorylation of eIF2α in chronic myeloid leukemia cells stimulates secretion of matrix modifying enzymes. Oncotarget 7, 79706–79721. 10.18632/oncotarget.12941

51. Podszywalow-Bartnicka, P., Kominek, A., Wolczyk, M., Kolba, M.D., Swatler, J., Piwocka, K., 2018. Characteristics of live parameters of the HS-5 human bone marrow stromal cell line cocultured with the leukemia cells in hypoxia, for the studies of leukemia-stroma cross-talk: Monitoring Stromal Cells in the Leukemia Coculture. Cytometry 93, 929–940. 10.1002/cyto.a.23580

52. Podszywalow-Bartnicka, P., Maifrede, S., Le, B.V., Nieborowska-Skorska, M., Piwocka, K., Skorski, T., 2019. PARP1 inhibitor eliminated imatinib-refractory chronic myeloid leukemia cells in bone marrow microenvironment conditions. Leukemia & Lymphoma 60, 262–264. 10.1080/10428194.2018.1471602

53. Rojek, K.O., Ćwiklińska, M., Kuczak, J., Guzowski, J., 2022. Microfluidic Formulation of Topological Hydrogels for Microtissue Engineering. Chem. Rev. 122, 16839–16909. 10.1021/acs.chemrev.1c00798

54. Saito, K., Zhang, Q., Yang, H., Yamatani, K., Ai, T., Ruvolo, V., Baran, N., Cai, T., Ma, H., Jacamo, R., Kuruvilla, V., Imoto, J., Kinjo, S., Ikeo, K., Moriya, K., Suzuki, K., Miida, T., Kim, Y.-M., Vellano, C.P., Andreeff, M., Marszalek, J.R., Tabe, Y., Konopleva, M., 2021. Exogenous mitochondrial transfer and endogenous mitochondrial fission facilitate AML resistance to OxPhos inhibition. Blood Advances 5, 4233–4255. 10.1182/bloodadvances.2020003661

55. Sart, S., Ronteix, G., Jain, S., Amselem, G., Baroud, C.N., 2022. Cell Culture in Microfluidic Droplets. Chem. Rev. 122, 7061–7096. 10.1021/acs.chemrev.1c00666

56. Schepers, K., Pietras, E.M., Reynaud, D., Flach, J., Binnewies, M., Garg, T., Wagers, A.J., Hsiao, E.C., Passegué, E., 2013. Myeloproliferative Neoplasia Remodels the Endosteal Bone Marrow Niche into a Self-Reinforcing Leukemic Niche. Cell Stem Cell 13, 285–299. 10.1016/j.stem.2013.06.009

57. Scielzo, C., Ghia, P., 2020. Modeling the Leukemia Microenviroment In Vitro. Front. Oncol. 10, 607608. 10.3389/fonc.2020.607608

58. Sharipol, A., Lesch, M.L., Soto, C.A., Frisch, B.J., 2022. Bone Marrow Microenvironment-On-Chip for Culture of Functional Hematopoietic Stem Cells. Front. Bioeng. Biotechnol. 10, 855777. 10.3389/fbioe.2022.855777

59. Simioni, C., Conti, I., Varano, G., Brenna, C., Costanzi, E., Neri, L.M., 2021. The Complexity of the Tumor Microenvironment and Its Role in Acute Lymphoblastic Leukemia: Implications for Therapies. Front. Oncol. 11, 673506. 10.3389/fonc.2021.673506

60. Sugiura, S., Nakajima, M., Iwamoto, S., Seki, M., 2001. Interfacial Tension Driven Monodispersed Droplet Formation from Microfabricated Channel Array. Langmuir 17, 5562–5566. 10.1021/la010342y

61. Swatler, J., Lo Tartaro, D., Borella, R., Brewinska-Olchowik, M., Paolini, A., Neroni, A., Turos-Korgul, L., Wiech, M., Kozlowska, E., Cysewski, D., Grabowska-Pyrzewicz, W., Wojda, U., Basak, G., Argüello, R.J., Cossarizza, A., De Biasi, S., Piwocka, K., 2022a. Dysfunctional subsets of CD39+ T cells, distinct from PD-1+, driven by leukemic extracellular vesicles in myeloid leukemias. haematol 108, 909–916. 10.3324/haematol.2022.281713

62. Swatler, J., Turos-Korgul, L., Brewinska-Olchowik, M., De Biasi, S., Dudka, W., Le, B.V., Kominek, A., Cyranowski, S., Pilanc, P., Mohammadi, E., Cysewski, D., Kozlowska, E., Grabowska-Pyrzewicz, W., Wojda, U., Basak, G., Mieczkowski, J., Skorski, T., Cossarizza, A., Piwocka, K., 2022b. 4-1BBL–containing leukemic extracellular vesicles promote immunosuppressive effector regulatory T cells. Blood Advances 6, 1879–1894. 10.1182/bloodadvances.2021006195

63. Tan, J.J.Y., Nguyen, D.-V., Common, J.E., Wu, C., Ho, P.C.L., Kang, L., 2021. Investigating PEGDA and GelMA Microgel Models for Sustained 3D Heterotypic Dermal Papilla and Keratinocyte Co-Cultures. IJMS 22, 2143. 10.3390/ijms22042143

64. Tobin, L.A., Robert, C., Rapoport, A.P., Gojo, I., Baer, M.R., Tomkinson, A.E., Rassool, F.V., 2013. Targeting abnormal DNA double-strand break repair in tyrosine kinase inhibitor-resistant chronic myeloid leukemias. Oncogene 32, 1784–1793. 10.1038/onc.2012.203

65. Velasco-Rodriguez, B., Diaz-Vidal, T., Rosales-Rivera, L.C., García-González, C.A., Alvarez-Lorenzo, C., Al-Modlej, A., Domínguez-Arca, V., Prieto, G., Barbosa, S., Soltero Martínez, J.F.A., Taboada, P., 2021. Hybrid Methacrylated Gelatin and Hyaluronic Acid Hydrogel Scaffolds. Preparation and Systematic Characterization for Prospective Tissue Engineering Applications. IJMS 22, 6758. 10.3390/ijms22136758

66. Vianello, F., Villanova, F., Tisato, V., Lymperi, S., Ho, K.-K., Gomes, A.R., Marin, D., Bonnet, D., Apperley, J., Lam, E.W.-F., Dazzi, F., 2010. Bone marrow mesenchymal stromal cells non-selectively protect chronic myeloid leukemia cells from imatinib-induced apoptosis via the CXCR4/CXCL12 axis. Haematologica 95, 1081–1089. 10.3324/haematol.2009.017178

67. Wang, H., Liu, Haitao, Liu, Hui, Su, W., Chen, W., Qin, J., 2019. One-Step Generation of Core–Shell Gelatin Methacrylate (GelMA) Microgels Using a Droplet Microfluidic System. Adv Materials Technologies 4, 1800632. 10.1002/admt.201800632

68. Wang, H., Zhou, L., Liao, J., Tan, Y., Ouyang, K., Ning, C., Ni, G., Tan, G., 2014. Cell-laden photocrosslinked GelMA–DexMA copolymer hydrogels with tunable mechanical properties for tissue engineering. J Mater Sci: Mater Med 25, 2173–2183. 10.1007/s10856-014-5261-x

69. Wang, K., Nune, K.C., Misra, R.D.K., 2016. The functional response of alginate-gelatin-nanocrystalline cellulose injectable hydrogels toward delivery of cells and bioactive molecules. Acta Biomaterialia 36, 143–151. 10.1016/j.actbio.2016.03.016

70. Wei, X., Huang, B., Chen, K., Fan, Z., Wang, L., Xu, M., 2022. Dot extrusion bioprinting of spatially controlled heterogenous tumor models. Materials & Design 223, 111152. 10.1016/j.matdes.2022.111152

71. Weisberg, E., Wright, R.D., McMillin, D.W., Mitsiades, C., Ray, A., Barrett, R., Adamia, S., Stone, R., Galinsky, I., Kung, A.L., Griffin, J.D., 2008. Stromal-mediated protection of tyrosine kinase inhibitor-treated BCR-ABL-expressing leukemia cells. Molecular Cancer Therapeutics 7, 1121– 1129. 10.1158/1535-7163.MCT-07-2331

72. Wolczyk, M., Serwa, R., Kominek, A., Klejman, A., Milek, J., Chwałek, M., Turos-Korgul, L., Charzyńska, A., Dabrowski, M., Dziembowska, M., Skorski, T., Piwocka, K., Podszywalow-Bartnicka, P., 2023. TIAR and FMRP shape pro-survival nascent proteome of leukemia cells in the bone marrow microenvironment. iScience 26, 106543. 10.1016/j.isci.2023.106543

73. Yang, W., Cai, S., Chen, Y., Liang, W., Lai, Y., Yu, H., Wang, Y., Liu, L., 2020. Modular and Customized Fabrication of 3D Functional Microgels for Bottom-Up Tissue Engineering and Drug Screening. Adv Materials Technologies 5, 1900847. 10.1002/admt.201900847

74. Yue, K., Trujillo-de Santiago, G., Alvarez, M.M., Tamayol, A., Annabi, N., Khademhosseini, A., 2015. Synthesis, properties, and biomedical applications of gelatin methacryloyl (GelMA) hydrogels. Biomaterials 73, 254–271. 10.1016/j.biomaterials.2015.08.045

75. Zhang, B., Chu, S., Agarwal, P., Campbell, V.L., Hopcroft, L., Jørgensen, H.G., Lin, A., Gaal, K., Holyoake, T.L., Bhatia, R., 2016. Inhibition of interleukin-1 signaling enhances elimination of tyrosine kinase inhibitor–treated CML stem cells. Blood 128, 2671–2682. 10.1182/blood-2015-11-679928

76. Zhang, B., Li, M., McDonald, T., Holyoake, T.L., Moon, R.T., Campana, D., Shultz, L., Bhatia, R., 2013. Microenvironmental protection of CML stem and progenitor cells from tyrosine kinase inhibitors through N-cadherin and Wnt–β-catenin signaling. Blood 121, 1824–1838. 10.1182/blood-2012-02-412890

77. Zhang, T., Zhang, H., Zhou, W., Jiang, K., Liu, C., Wang, R., Zhou, Y., Zhang, Z., Mei, Q., Dong, W.-F., Sun, M., Li, H., 2022. One-Step Generation and Purification of Cell-Encapsulated Hydrogel Microsphere With an Easily Assembled Microfluidic Device. Front. Bioeng. Biotechnol. 9, 816089. 10.3389/fbioe.2021.816089

78. Zhao, X., Lang, Q., Yildirimer, L., Lin, Z.Y., Cui, W., Annabi, N., Ng, K.W., Dokmeci, M.R., Ghaemmaghami, A.M., Khademhosseini, A., 2016a. Photocrosslinkable Gelatin Hydrogel for Epidermal Tissue Engineering. Adv. Healthcare Mater. 5, 108–118. 10.1002/adhm.201500005

79. Zhao, X., Liu, S., Yildirimer, L., Zhao, H., Ding, R., Wang, H., Cui, W., Weitz, D., 2016b. Injectable Stem Cell-Laden Photocrosslinkable Microspheres Fabricated Using Microfluidics for Rapid Generation of Osteogenic Tissue Constructs. Adv Funct Materials 26, 2809–2819. 10.1002/adfm.201504943

80. Zippel, S., Dilger, N., Chatterjee, C., Raic, A., Brenner-Weiß, G., Schadzek, P., Rapp, B.E., Lee-Thedieck, C., 2022. A parallelized, perfused 3D triculture model of leukemia for in vitro drug testing of chemotherapeutics. Biofabrication 14, 035011. 10.1088/1758-5090/ac6a7e

