## Supplementary Figures for "High-throughput formulation of reproducible 3D cancer microenvironments for drug testing in myelogenous leukemia"

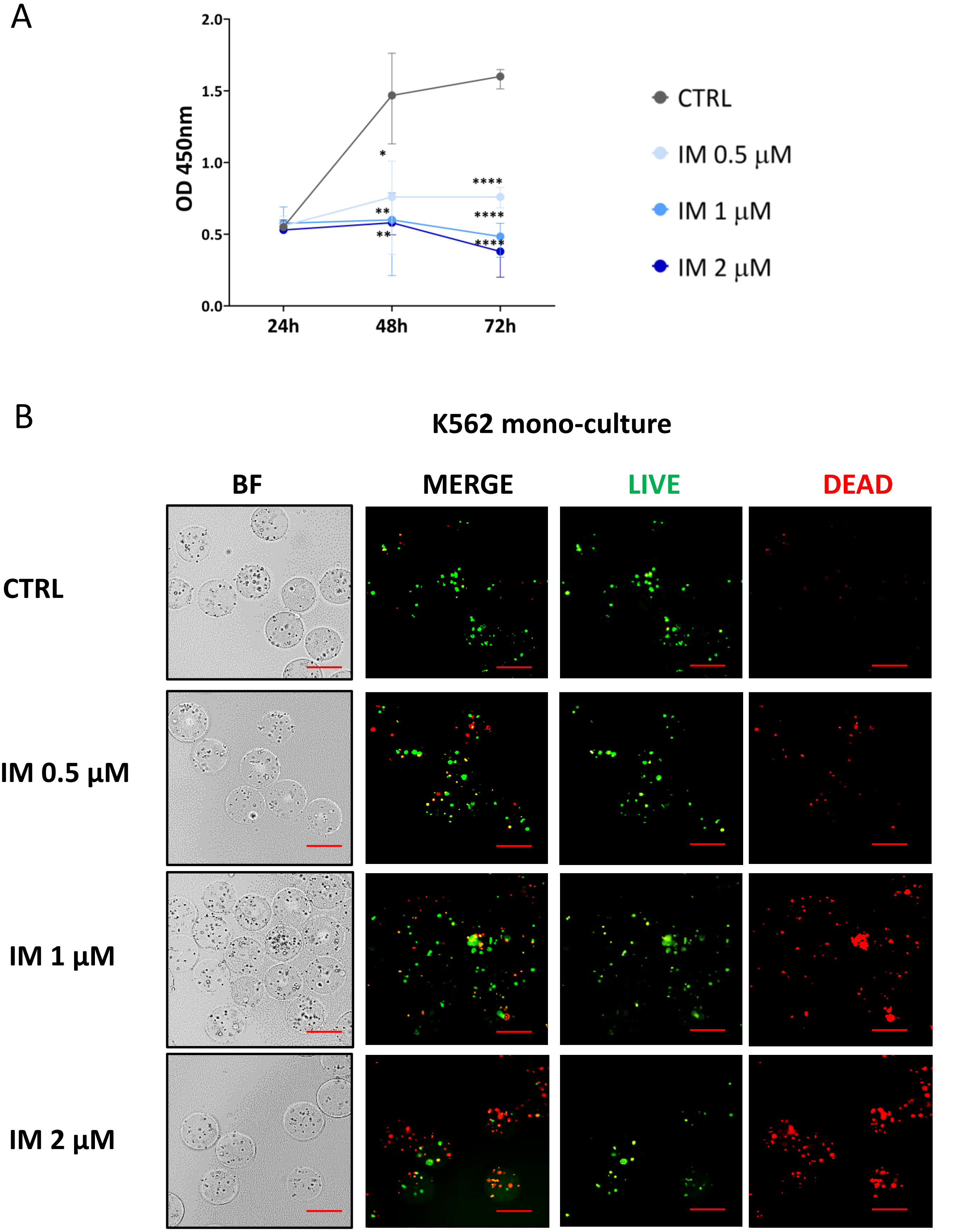


**Supplementary Figure S1. (A)** CCK-8 results for imatinib treatment of cancer K562 monocultures encapsulated in 4.8% GelMA microbeads. **(B)** The corresponding LIVE/DEAD staining (green/red). One can observe increased number of dead cells upon imatinib treatment as compared to control (no drug).

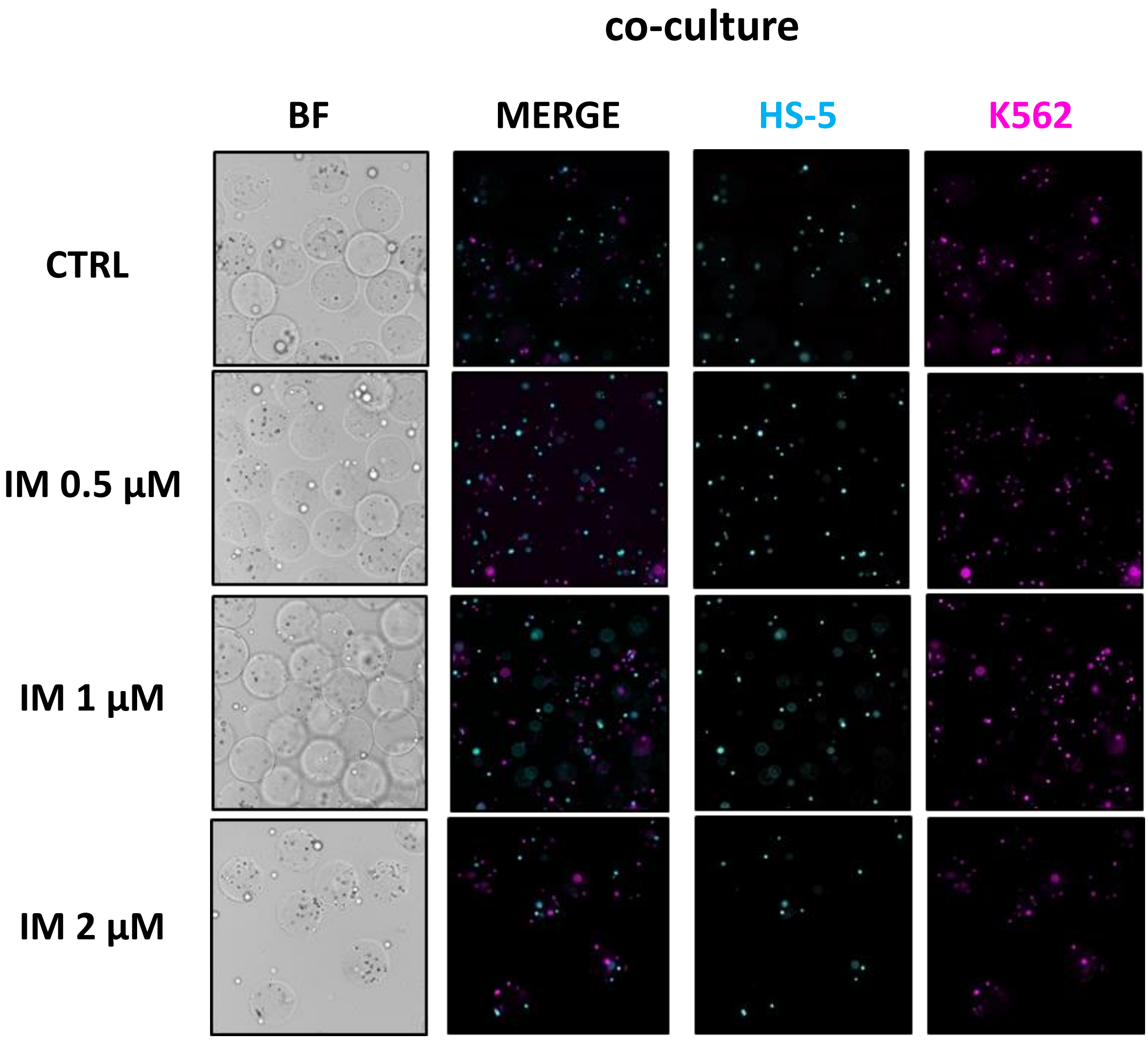


**Supplementary Figure S2.** Fluorescently labelled K562 cells (magenta) and HS-5 cells (cyan) encapsulated in 4.8% GelMA microbeads, imaged 48 h post encapsulation upon Imatinib treatment. The cells were labeled with CellTracker™ Red CMTPX Dye (K562) and CellTracker™ Green CMFDA Dye (HS-cells) and re-colored upon image postprocessing. One can observe increased magenta signal which may indicate faster growth of cancer cells as compared to stroma upon Imatinib treatment. Pictures were taken using fluorescence Nikon microscope, scale bar = 200 µm.

A

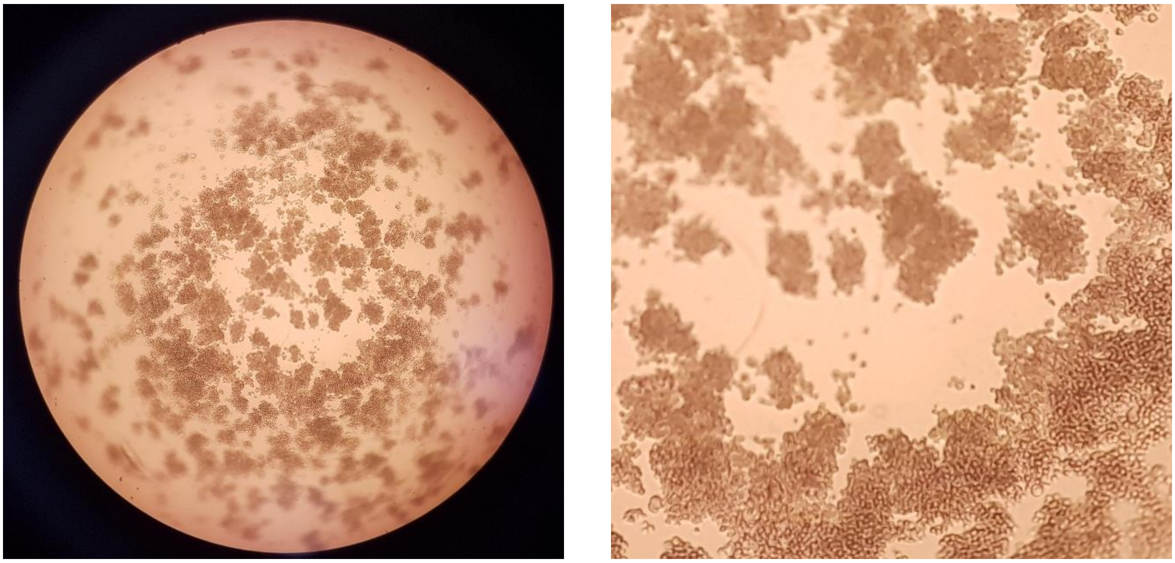

B

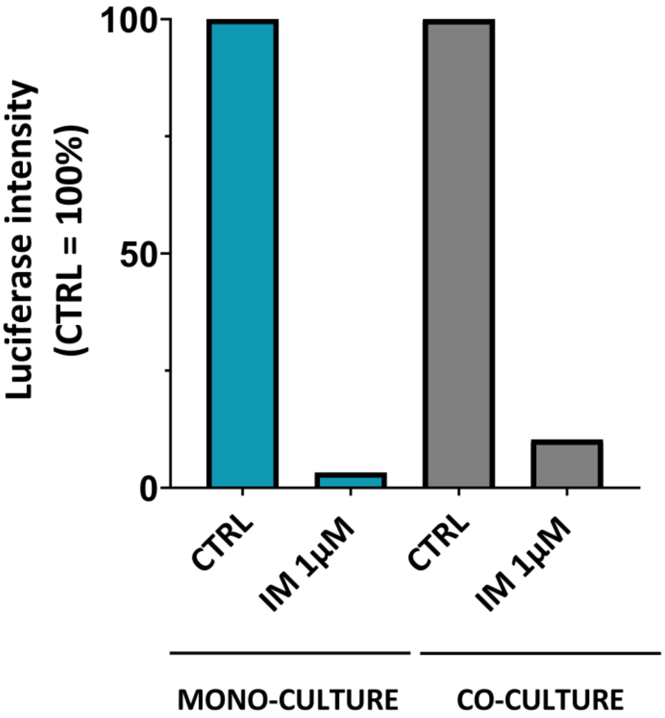


**Supplementary Figure S3.** Macroencapsulation of K562 monoculture and co-culture (1:1) of K562 and HS-5 cells in 5% matrigel under RPMI medium supplemented with L-glutamine. K562 cells were lentivirally transduced with plasmid pLenti7.3-redluc expressing GFP and Firefly luciferase, followed by flow cytometry cell sorting of GFP-positive cells. After a week of **culture**, 1uM imatinib was added for 72h. **(A)** Brightfield imaging of microtissues formed in co-culture using Motic AE20 microscope with objective 4x. **(B)** Leukemic cells growth after imatinib treatment assessed as activity of luciferase in the cell lysates compared to control (untreated) = 100%.
